# Kappa opioid receptor activation induces epigenetic silencing of brain-derived neurotropic factor via HDAC5 in depression

**DOI:** 10.1101/2023.09.18.558045

**Authors:** Anubhav Yadav, Shalini Dogra, Boda Arun Kumar, Poonam Kumari, Ajeet Kumar, Manish K Dash, Prem N Yadav

## Abstract

Treatment-resistant depression (TRD) occurs in almost 50% of depressed patients. Central kappa opioid receptor (KOR) agonism has been demonstrated to induce depression and anxiety, while KOR antagonism alleviate depression like symptoms in rodent models and TRD in clinical studies. Previously, we have shown that sustained KOR activation leads to TRD-like phenotype in mice, and modulation of brain-derived neurotrophic factor (BDNF) expression in the prefrontal cortex (PFC) appears to be one of the molecular determinants of the antidepressant response. In the present study, we observed that sustained KOR activation by a selective agonist, U50488, selectively reduced the *Bdnf* transcripts *II, IV*, and *Bdnf CDS* (protein-coding *Exon IX*) in the PFC and cultured primary cortical neurons, which was blocked by selective KOR antagonist, norbinaltorphimine. Considering the crucial role of epigenetic pathways in *BDNF* expression, we further investigated the role of various epigenetic markers in KOR induced BDNF downregulation in mice. We observed that treatment with U50488 resulted in selective and specific downregulation of acetylation at the 9th lysine residue of the histone H3 protein (H3K9ac) and upregulation of HDAC5 expression in the PFC. Further, using anti-H3K9ac and anti-HDAC5 antibodies in chromatin immune precipitation assay, we detected decreased enrichment of H3K9ac and increased HDAC5 binding at *Bdnf II* and *IV* transcripts after U50488 treatment, which were blocked by a selective KOR antagonist, norbinaltorphimine. Further mechanistic studies using HDAC5 selective inhibitor, LMK235, in primary cortical neurons, and adeno-associated viral shRNA mediated HDAC5-knockdown in the PFC of mice, we demonstrated an essential role of HDAC5 in KOR-mediated reduction of *Bdnf* expression in the PFC and depression-like symptoms in mice. These results suggest that KOR engages multiple pathways to induce depression-like symptoms in mice, and provide novel insights into the mechanisms by which activation of KOR regulates major depressive disorders.

## Introduction

As per WHO estimates (March 2023), about 3.8% of the world population is affected by depressive disorders (https://www.who.int/news-room/fact-sheets/detail/depression), and notably, almost 50% of these do not respond to the current therapeutic interventions leading to a treatment-resistant depression (TRD) condition[1]. Therefore, TRD is a highly unmet medical need, and the neurobiological distinction between treatment resistant and treatment-responsive depression are not clear. Although esketamine was approved for TRD in 2019 and major depressive disorders with suicidal ideation in 2020 [2], it causes serious adverse effects such as blood pressure increase, vertigo, and dissociation [3]. Several mechanistic studies in rodent models of depression suggest that salutary effects of esketamine are mediated via regulation of brain-derived neurotrophic factor (BDNF) expression, synthesis, and release, which in turn activate synaptic plasticity genes [4, 5]. Further, decreased BDNF expression is also considered as key factors responsible for impaired synaptic plasticity observed in the rodent models of depression and in depressed patients [6]. BDNF is critical for brain development and several neuro-physiological processes, and is highly conserved among all mammalian species. The expression and function of BDNF are tightly regulated by eight exons with individual promoters that undergo tissue- and stimulus-specific alternative splicing to generate multiple *Bdnf* transcripts with distinct 5’ non-coding exons as a promoter with the constitutive *exon IXa* (protein coding) gives rise to different Bdnf transcripts [7–9]. Interestingly, various stressors are known to change the levels of *Bdnf* transcripts in different brain regions, and treatment with antidepressants can restore it [10–12], but mechanisms of the regions-specific regulation of various *Bdnf* transcripts in TRD are not clear, yet.

Kappa opioid receptor (KOR) activation in mesolimbic circuit reciprocally modulates mood via inhibiting ventral tegmental area dopaminergic neurons firing. This decrease in dopaminergic neurotransmission KOR activation is pivotal to dysphoria and anhedonia-related symptoms, as well as the modulation of mood and stress [13, 14]. Recent clinical studies using KOR antagonist, CERC501, or a combination of buprenorphine and samidorphan in patients suffering from major depressive disorders (MDD) have demonstrated KOR as a promising target for TRD [15, 16]. We have also shown that chronic activation of KOR induces TRD-like phenotype in mice, which are reversed by pre-treatment with KOR selective antagonist, norbinaltorphimine (norBNI), but not by selective serotonin reuptake inhibitors (SSRIs), fluoxetine and citalopram [17]. In this study, we also observed that KOR activation led to decreased BDNF expression in the prefrontal cortex (PFC) that was blocked by norBNI, but not by Fluoxetine or Citalopram or Imipramine, suggesting BDNF levels in PFC as molecular determinants of antidepressant response. Although involvement of epigenetic modifications and histone deacetylases (HDACs) are widely known to regulate BDNF expression in depressive disorders [12, 18, 19], the mechanisms responsible for KOR-Induced downregulation of BDNF expression in depression are not known, yet. Therefore, in the present study, we sought to investigate the effect and mechanism of chronic KOR activation on the expression of major *Bdnf* transcripts in the PFC and primary cortical neurons. We observed that sustained KOR activation selectively reduced the *Bdnf* transcripts *II, IV*, and *Bdnf CDS* (protein-coding *Exon IX*), which was blocked by selective KOR antagonist, norBNI. Further, mechanistic studies in mice and primary cortical neurons revealed that chronic KOR activation lead to decreased acetylation at the 9th lysine residue of the histone H3 protein (H3K9ac) and upregulation of HDAC5 binding at *Bdnf II* promoter, which suppresses the *Bdnf* expression in the PFC and ultimately depression like symptoms in mice.

## Material and Methods

### Animals

*In vivo* experiments and procedures were performed according to the guidelines provided by the Institutional Animal Ethics Committee (IAEC) of CSIR-Central Drug Research Institute, Lucknow, India. C57BL/6J male mice of 6-8 weeks and weight 22-25 g were used in this study. Mice were housed in a 12-h light/dark cycle under constant temperature (22-25°C) and consistent humidity (50 ± 5%) with *ad libitum* food and water supply.

### Drugs

KOR agonist (−)-*trans*-(1S,2S)-U-50488 hydrochloride hydrate (Sigma Aldrich, U111) (U50488) was administered intraperitoneally (*i.p.*) at a dose of 5 mg/kg for 21 days. The KOR selective antagonist, *nor*-Binaltorphimine dihydrochloride (norBNI; Sigma Aldrich, N1771, 10mg/kg, *i.p.*) was administered only once on day 1, 30 minutes before the U50488 treatment. Control animals were injected with vehicle solution containing 0.9% sodium chloride+0.1% DMSO.

### Primary neuronal culture

Primary neurons were cultured from the cerebral cortices of 0-1 day-old mouse pups as described previously [17]. Briefly, the cortices were gently dissociated in 1X Hanks′ Balanced Salt Solution (HBSS) (Sigma Aldrich-# H4641) and were digested with papain (0.1% Sigma Aldrich-#P4762) for 20 minutes at 37 °C. After digestion, tissue samples were triturated 10-15 times to get the dissociated cells, which were then suspended in neurobasal media (Gibco, #12348-017) supplemented with 1X B27 supplement (Invitrogen, #17504) and 0.5mM Glutamax (Gibco, #35050). The cell suspension was then plated onto a 12-well tissue culture plate (0.5×10^6^ cells/well) coated with Poly-L-lysine (PLL) (0.1mg/ ml Sigma Aldrich, # 2636P) and incubated at 37°C in a humidified atmosphere of 95% oxygen and 5% CO_2_.

### Reverse transcription (RT) and quantitative PCR

Mice were euthanized after 24 h of last administration of U50488 by the overdose of an anesthetic avertin (350 mg/kg). Brain tissues - prefrontal cortex (PFC), striatum, and hippocampus were rapidly micro-dissected after the decapitation of fully anesthetized mice and immediately snap-frozen in liquid nitrogen and stored at -80°C till further use. Total RNA was extracted from the brain tissues using the trizol/chloroform method. First, total RNA was treated with DNAase at 37°C for 10 min to remove genomic DNA, then reverse transcribed into cDNA (iScript™ cDNA Synthesis Kit, #1708891). This cDNA was then used for real-time PCR detection using the exon-specific primers (**Supplementary Table 1**). The condition for PCR was 95°C for 10 minutes, followed by 40 cycles of 95°C for 15s, 56°C for 30s. The housekeeping gene β-actin or GAPDH was used as a reference gene for the normalization of gene expression. The 2−ΔΔCTmethod, (i.e., delta-delta-ct analysis) was used for relative quantification (Valasek and Repa, 2005).

### Western blotting

Western blotting was used to determine the expression of various protein and was performed as described earlier [17]. Briefly, frozen brain tissues were homogenized in lysis buffer (25 mM Tris-HCl, pH 7.6,150 mM NaCl, 0.5% NP-40,1% sodium deoxycholate,0.1% SDS). Briefly, 40-50 µg of protein samples were subjected to 8-12% SDS-PAGE and transferred to a polyvinyl difluoride membrane followed by blocking for two hours at room temperature by blocking buffer (5% non-fat dry milk and 3% bovine serum albumin in PBST). Specific primary antibodies (**Table S2**) were incubated overnight at 4°C, followed by incubation with horseradish peroxidase conjugated secondary antibodies (1:4000) at room temperature for 2 hours. Blots were developed by enhanced chemiluminescence substrate and images were acquired using Amersham™ ImageQuant™ 800 and densitometric analysis was performed by ImageJ software.

### Immunohistochemistry

C57BL/6J mice were transcardially perfused with a 4% paraformaldehyde (PFA) solution. The brains were removed, post-fixed overnight in 4% PFA, and dehydrated in 30% sucrose for 24 hours. Thin coronal slices (30 μm) were made using a cryostat (FSE, ThermoScientific). For staining, the sections were with 1 X phosphate buffer saline (PBS0. The antigen retrieval was performed by boiling slices in 10 mM sodium citrate buffer (pH-6.3) for 20 minutes. Slices were incubated with 0.5% TritonX 100 solution (in 1X PBS) for one hour at room temperature followed by blocking (3% bovine serum albumin, 3% horse serum, 0.3% Triton X-100 in 1X PBS) for two hours at room temperature. The sections were then incubated with mouse anti-HDAC5 (1:50) overnight at 4°C. After overnight incubation, the brain sections were washed three times with PBST buffer (0.1% Triton X 100 in 1X PBS) followed by incubation with respective secondary antibodies conjugated with Alexa fluor 594 or 488 (1:500) and Hoechst 33258 (2 µg/ml) in blocking buffer for 1 hour at room temperature. After incubation with host specific secondary antibodies, the slices were washed thrice and were mounted onto poly-lysine-coated glass slides using Vectashield antifade mounting medium, and fluorescent images were collected with XYZ acquisition mode at 1μm step size using the Olympus BX61-FV1200-MPE microscope using a 40X oil (1.3NA) objective at 8.0μs/pixel scan speed. (Olympus, Shinjuku, Tokyo, Japan). Pixel intensity quantification for HDAC5 and Hoechst 33258 staining in anterior cingulate cortex and piriform cortex was performed using ImageJ software (www.rcb.info.nih.gov/ij) and presented as fold change after normalization with pixel intensity of Hoechst 33258 in each region.

### Chromatin immunoprecipitation (ChIP)

ChIP was performed as described earlier [20] with slight modifications. Freshly dissected PFC tissues were crosslinked using 1% formaldehyde in 1XPBS for 10 minutes at ice, followed by quenching with 125 mM glycine for 10 minutes. Cross linked PFCs were then washed with 1X PBS and homogenized in a SDS lysis buffer (0.01% SDS, 100 mM EDTA, 1 mM PMSF, 50 mM Tris-HCl, pH 8.1) and sonicated on ice (FB-705; Fisher Scientific) at 40% amplitude and a 10 × 10 second pulse. Sonicated samples were spun at 12,000 rcf for 10 minutes at 4°C, and supernatants were collected for dilution in the ChIP dilution buffer (167 mM NaCl, 0.01% SDS, 1.1% Triton X-100, 1.2 mM EDTA, 16.7 mM Tris-HCl, pH 8.1) and 10% of this DNA sample was kept aside as input DNA for qPCR and fold change calculations at later stage. The supernatant containing sheared chromatin (50 µg) was incubated with either nonimmune immunoglobulin (IgG) (as a control) or anti-H3K9ac or anti-HDAC5 anitbodies for 14 hours at 4°C followed by centrifugation for 2 minutes at 3000 rcf at 4°C. DNA-histone complex was collected by binding to salmon sperm DNA blocked-protein A-agarose beads for a 3-4 hours incubation at 4°C and then sequentially washed with low salt buffer (150 mM NaCl, 0.1% SDS, 1% Triton X-100, 2 mM EDTA, 20 mM Tris-HCl, pH 8.1), high salt buffer (500 mM NaCl, 0.1% SDS, 1% Triton X-100, 2 mM EDTA, 20 mM Tris-HCl, pH 8.1), LiCl immune complex buffer (0.25M LiCl, 1% IGEPAL-CA630, 1% deoxycholic acid, 1 mM EDTA, 10 mM Tris, pH 8.1) and Tris-EDTA buffer (10 mM Tris-HCl, 1 mM EDTA, pH 8.0). DNA-histone complexes were removed from protein A-agarose beads by incubation with elution buffer (0.1 M NaHCO3, 1% SDS) for 20 minutes at room temperature. The eluted DNA-histone complexes were then digested with proteinase K at 55°C for overnight followed by DNA extraction using the phenol-chloroform-alcohol method. An equal volume of immunoprecipitated DNA and input DNA was subjected to quantitative real-time PCR using mouse *Bdnf exon* specific primers.

### HDAC5 knockdown studies in mice

A casset of U6 promoter followed by mouse HDAC5 specific shRNA [21] was synthezised and cloned between Bsu36I and SpeI site of adeno associated viral (AAV) vector pTR-CBA-TdTomato and packaged into AAV9 serotype using pXX6-80 and pXR9 (kind gift from Dr. Aravind Asokan, UNC-Chapel Hill, USA) in HEK293 as described earlier [17]. Briefly, HDAC5-shRNA virons were purified using a discontinuous iodixanol gradient (Sigma-Aldrich) and concentrated to 1 × 10^12^ genome copy/ml, and 0.5 µl of these viral particles were delivered bilaterally into the anterior singulate cortices (ACC, AP: -1 mm, ML: ± 0.2 mm, DV: 1.5 mm) bilaterly using a motorized stereotaxic system (Stoelting, USA). Sterotaxic surgeries were performed using a mixture of ketamine and xylazine (90 and 10 m/ kg, respectively) as anaesthetic agents and Meloxicam (1 mg/ kg, three consecutive days, *s.c*.) as a postoperative analgesia. A knockdown efficiency was measured by quantitative (qPCR) using RNA isolated from PFCs after completion of the experiment. Further, the region specificity of the viral expression was verified in each mouse using immunohistochemistry. Mice expressing inadequate viral expression were excluded from the analysis.

### Neurobehavioral studies

#### Forced swim test

Forced swim test was performed to determine the behavioural despair as described previously (Dogra et al, 2016). Briefly, mice were placed into a clear Plexiglass cylinder (25 cm in height and 10 cm in diameter) filled with water (25-27°C) to a depth of 20 cm. A 5 minutes’ swim session was video-recorded by a camera mounted on tripod alongside of the cylinder to capture the whole animal movements, including 4 limbs and tail. The duration of immobility was determined using AnyMaze7.1 software. Immobility was defined as the absence of all movements except those required for respiration.

#### Tail suspension test

Mice were subjected to a tail suspension test after 21 days of U50 treatment. The total duration of immobility induced by the tail suspension was measured according to the method. Mice were hung 50 cm above the floor by adhesive tape positioned roughly 2 cm from the tip of the tail. During a 5-minute test session, the time mice stayed motionless was measured. Mice were considered immobile only when they hung passively and motionless. AnyMaze7.1 software was used to record a 5-minute video and analysis of immobility time of the tail suspension test.

#### Social behavior

The animals’ sociability was assessed using a three-chambered social behaviour test box, as previously described [17]. Test mouse was permitted to explore the entire space for 10 minutes (acclimatisation phase). Following acclimatisation, a new mouse (C57BL/6J, stranger) was placed in a wired cage housed in one of the side chambers (social chamber). The test mouse was placed in the centre compartment and given 10 minutes to explore the all chambers. A camera was used to capture mouse behaviour, and the time spent in each chamber was analysed using AnyMaze 7.1 software. Sociability index was calculated as (time in social chamber-time spent in empty chamber)/ total time spent in both chamber.

## Results

### 1. Effect of chronic KOR activation on the expression of various *Bdnf* transcripts in the prefrontal cortex

In our previous study, we demonstrated that chronic treatment of U50488 (5mg/kg, *i.p*., 21 days) develops a treatment-resistant depression like phenotype. We have also shown that treatment with U50488 decreased *Bdnf* expression in the prefrontal cortex (PFC) and the hippocampus of mice. Interestingly, imipramine, fluoxetine, and citalopram attenuated U50488-induced decreases in the *Bdnf* expression in the hippocampus, but not in the PFC, indicating a critical role of the PFC BDNF signaling in antidepressant response observed in the chronic U50488 treated mice [17]. In the present study, we investigated underlying mechanisms of KOR-induced downregulation of BDNF protein expression in the PFC. We used previously reported U50488 administration regimen (5 mg/kg; *i.p*., 21 days) and immobility time in the forced swim test as a surrogate measure of depressive behavior [17]. As expected, we observed a significant increase in the immobility time after chronic treatment with selective KOR agonist, U50488 (**Figure S1).** We then determined if KOR activation-induced decreases in BDNF protein expression in the PFC is regulated by alternative usage of five major non-coding *5’exons* (*exon I-V*) acting as promoters for generating identical protein encoded by *exon IXa,* referred a*s Bdnf CDS* [7]. Indeed, we observed that chronic treatment with U50488 significantly decreased levels of *Bdnf II, IV*, and *Bdnf CDS* transcripts (**Figure 1A**), and increased the level of *Bdnf III* transcript (**Figure S2A**) as compared to the vehicle treated animals. A single dose of selective KOR antagonist, norBNI (10 mg/kg, *i.p*.) administered 30 minutes prior to the first U50488 treatment on day-1 blocked the effects of U50488 on *Bdnf* transcripts in the PFC (**Figure1A, Figure S2A**). Similar to *in vivo* effects of KOR activation, we also observed U50488 (100 nM and 1000 nM, for 72 hours) treatment-induced reduction in the levels of *Bdnf II*, IV, and *CDS* transcripts (**Figure 1B**) and increased the levels of *Bdnf III* transcript (**Figure S2B**) in the cultured primary cortical neurons. These results indicate that chronic KOR activation decreases BDNF protein expression in the PFC by inhibiting expression of *Bdnf II* and *IV* transcripts.

**Figure 1.**
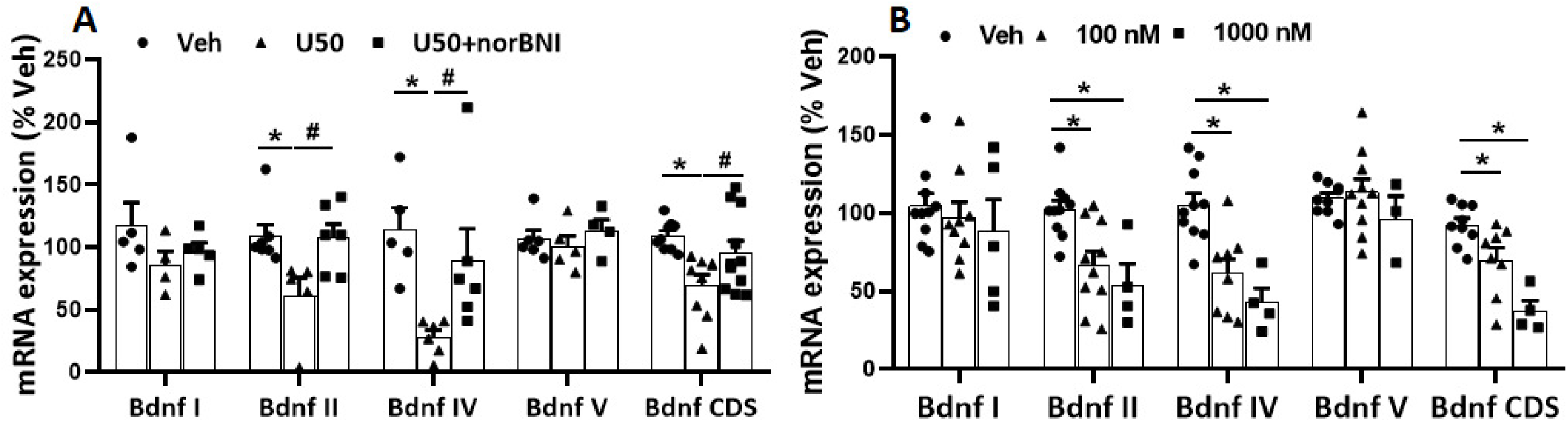
KOR activation leads to differential regulation of *Bdnf* transcripts in the prefrontal cortex (PFC). (**A**) Chronic treatment with U50488 (U50; 5 mg/kg *i.p*.; 21 days) significantly decreased the levels of *Bdnf II, Bdnf IV*, and *Bdnf CDS (*protein coding *exon IXa)* transcripts in the prefrontal cortex (PFC). A single administration of KOR antagonist, *nor*-binaltorphimine (norBNI, *i.p*.; 10 mg/kg), blocked the effects of U50488 treatment on the expression of *Bdnf* transcripts. U50488 treatment did not change the levels of *Bdnf I* and *V* transcripts in the PFC. Data are presented as mean ± SEM of % Vehicle). *Bdnf II*: *p*= 0.018, *F*_(2,15)_= 5.284; *Bdnf IV*: *p*= 0.014, *F*_(2,14)_= 5.784; *Bdnf CDS: p*= 0.0087, *F*_(2,26)_= 5.721), \**p*<0.05 (Veh vs U50), ^#^*p*<0.05 (U50 vs U50+norBNI) by One way ANOVA followed by Newman-Keul’s multiple comparisons test, n= 5-7 mice/group. (**B**) Representative bar graph showing reduced expression of *Bdnf II, Bdnf IV,* and *Bdnf CDS* transcripts (normalized to GAPDH transcript) in 12 days *in vitro* (DIV12) primary cortical neurons treated with U50488 (100 nM and 1000 nM;72 hours). U50488 treatment did not change the levels of *Bdnf I* and *V* transcripts in the primary cortical neurons. Data are presented as mean ± SEM of % Vehicle. *Bdnf II*: *p= 0.009; *F*_(2,21)_= 5.835, *Bdnf IV*: \**p*= 0.0014; *F*_(2,19)_= 9.743, *Bdnf CDS:* \**p*<0.05; *F*_(2,18)_= 5.125 by One way ANOVA followed by Newman-Keul’s multiple comparisons test, n= 6-8 /group.

### 2. Chronic KOR activation induces epigenetic modifications in the PFC

The differential expression of *Bdnf II*, *III*, *IV* transcripts prompted us to investigate the underlying mechanisms of these changes induced by KOR activation. We first determined the effects of U50488 treatment on the PFC levels of post-translational histone modifications that are known to regulate gene expression in the central nervous system (CNS). Hence we evaluated the expression of two active chromatin markers, H3K9ac and H3K4me3, and two repressive chromatin markers, H3K9me3 and H3K27me3 in the PFC tissues after various treatments. We found that treatment with U50488 selectively reduced the levels of H3K9ac in the PFC as compared to the vehicle treatment (**Figure 2A, B**), while no effect was observed on the levels of H3K4me3, H3K9me3 and H3K27me3. We further tested the effect of KOR stimulation in the primary cortical neurons and, similar to our results in the PFC, we observed that treatment with selective KOR agonist, U50488 (100 nM for 72 hours), significantly decreased the expression of H3K9ac (**Figure 2C, D and S3A, B**). Pre-treatment with selective KOR antagonist, norBNI blocked the U50488-induced decreases in the levels of H3K9ac in the primary cortical neurons (**Figure 2C, D**). Similar to the PFC tissue, we found no changes in the levels H3K9me3 and H3K27me3 in the primary cortical neurons treated with U50488 for 72 hours (**Figure S3E, 3F, 3G, 3H**). We also observed no change in the levels of H3K4me3 after 24 hours of treatment with U50488 in the primary neurons, while there was a reduction in the levels of H3K4me3 at the 48 and 72hour time points (**Figure S3C and D**). These results indicate overall repressive effects of KOR activation on H3K9ac modification that may regulate histone structure and modulate transcription factor binding. Interestingly, however, we didn’t observe any significant change in the H3K9ac expression in C57BL/6J mice exposed to chronic unpredictable stress (CUS) paradigm (**Figure S4 A, B**), indicating that change in levels of H3K9ac could be a marker of treatment-resistant depression-like condition induced by chronically activated KORs. Therefore, we performed further studies to delineate the underlying mechanisms and the role of H3K9ac in KOR-induced suppression of *Bdnf* expression in the PFC. We performed a series of chromatin Immunoprecipitation (ChIP) assays to study the levels of H3K9ac binding at different *Bdnf* promoters in the PFC tissues. We found significantly lesser enrichment of H3K9ac modification at *Bdnf* II, IV, and V promoters in the PFC of U50488-treated mice as compared to the vehicle-treated mice (**Figure 2E**). These results suggest that chronic U50488 treatment differentially decreases the expression of an active chromatin marker, H3K9ac, at various *Bdnf* promoters which may lead to the decreases in the levels of specific *Bdnf* mRNA observed in the PFC.

**Figure 2.**
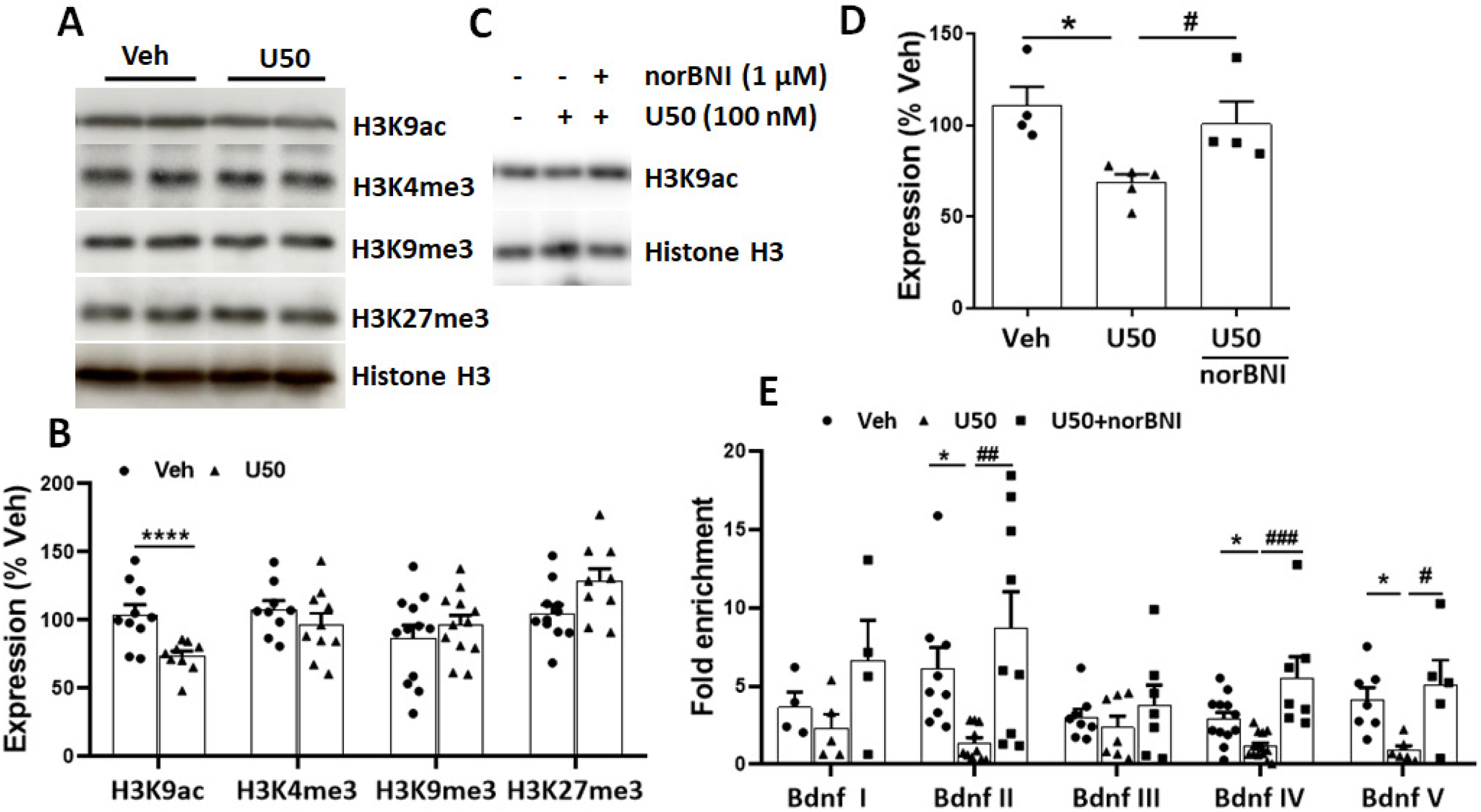
Chronic KOR activation causes epigenetic changes in the prefrontal cortex (PFC) of mice. (**A**) Chronic treatment with U50488 (5 mg/kg *i.p*.; 21 days) significantly decreased the expression of acetylation at the 9^th^ lysine residue of histone H3 (H3K9ac) in the PFC compared to the Vehicle (Veh)-treatment. No change in the levels of trimethylation at 4^th^, 9^th^ and 27^th^ lysine residues of histone H3 (H3K4me3, H3K9me3 and H3K27me3, respectively) was observed after treatment with U50488. (**B**) Bar graphs showing the quantification of the immunoblots presented in panel (A) (\*\*\*\**p*<0.0001 by unpaired t-test, t= 5.057, n= 6-8 mice/group). (**C**) Treatment of primary cortical neurons with U50488 (100 nM, 72 hours) significantly reduced the expression of H3K9ac, which was blocked by pre-treatment with a selective KOR antagonist, norBNI, (1 μM). (**D**) Bar graph showing the quantitative analysis of immunoblots presented in panel **C** (*p*= 0.0178, *F*_(2,10)_= 6.192), \**p*<0.05 (Veh vs U50), ^#^*p*<0.05 (U50 vs U50+*nor*BNI) by One-way ANOVA followed by Newman-Keul’s multiple comparisons test, n= 4-5/group. (**E**) Chromatin immunoprecipitation (ChIP) using anti-H3K9ac antibody showed a significant reduction in the H3K9ac enrichment at *Bdnf II, IV & V* promoters in the PFC. Pre-treatment with the KOR antagonist, norBNI, blocked the effects of U50488 treatment on H3K9ac binding at various *Bdnf* promoters in the PFC. The H3K9ac enrichment at *Bdnf I* and *III* promoters in the PFC did not differ between the treatment groups. *Bdnf II* (n= 9-11) *p*= 0.0066 *F*_(2,25)_= 6.184, *Bdnf IV* (n= 7-12); *p*= 0.0003 *F*_(2,33)_= 10.68, *Bdnf V* (n= 5-7) *p*= 0.0183 *F*_(2,15)_= 5.287 by One way ANOVA followed by Newman-Keul’s multiple comparisons test; \**p*<0.05, ^#^*p*<0.05, ^##^*p*<0.01, ^###^*p*<0.001. Data are presented as mean ± SEM of fold change over input.

### 3. KOR activation selectively modulates histone deacetylase 4 and 5 in the PFC

Considering that HDACs-induced deacetylation of lysine residues on histone tails regulates *Bdnf* expression [22, 23] and our results showing decreased levels of H3K9ac in the PFC of chronic U50488-treated mice, we sought to investigate the expression profile of various HDACs in the PFC after chronic treatment with U50488. To study this, we measured the protein levels of three of the four classes of HDACs, class 1 HDACs (HDAC1, HDAC2, HDAC3,), class 2 HDACs (HDAC4, HDAC5, HDAC6, HDAC7, HDAC8), and class 3 HDACs/Sirtuins (SIRT1-SIRT7) using western blotting. Among these HDACs, we found a significant decrease in the expression of HDAC4 and a significant increase in the expression of HDAC5 in the PFC of mice chronically treated with U50488, and both of these HDACs were normalized to control level by KOR antagonist norbinaltorphimine (**Figure 3A-D**, and **Figure S5A, B)**. Previously, chronic treatment with the antidepressant imipramine has also been shown to alleviate depression like behaviors by decreasing the expression of HDAC5 in the hippocampus, suggesting hippocampal levels of HDAC5 as a marker of antidepressant response [12]. Along that lines, we also profiled specific subregions of the PFC for U50488-induced changes in the levels of HDAC5 expression using immunohistochemistry. Our analysis of the PFC regions revealed that U50488 treatment selectively increased the expression of HDAC5 in the anterior cingulate (**Figure 4 and Figure S7**) and piriform cortex regions (**Figure S8).**

**Figure 3.**
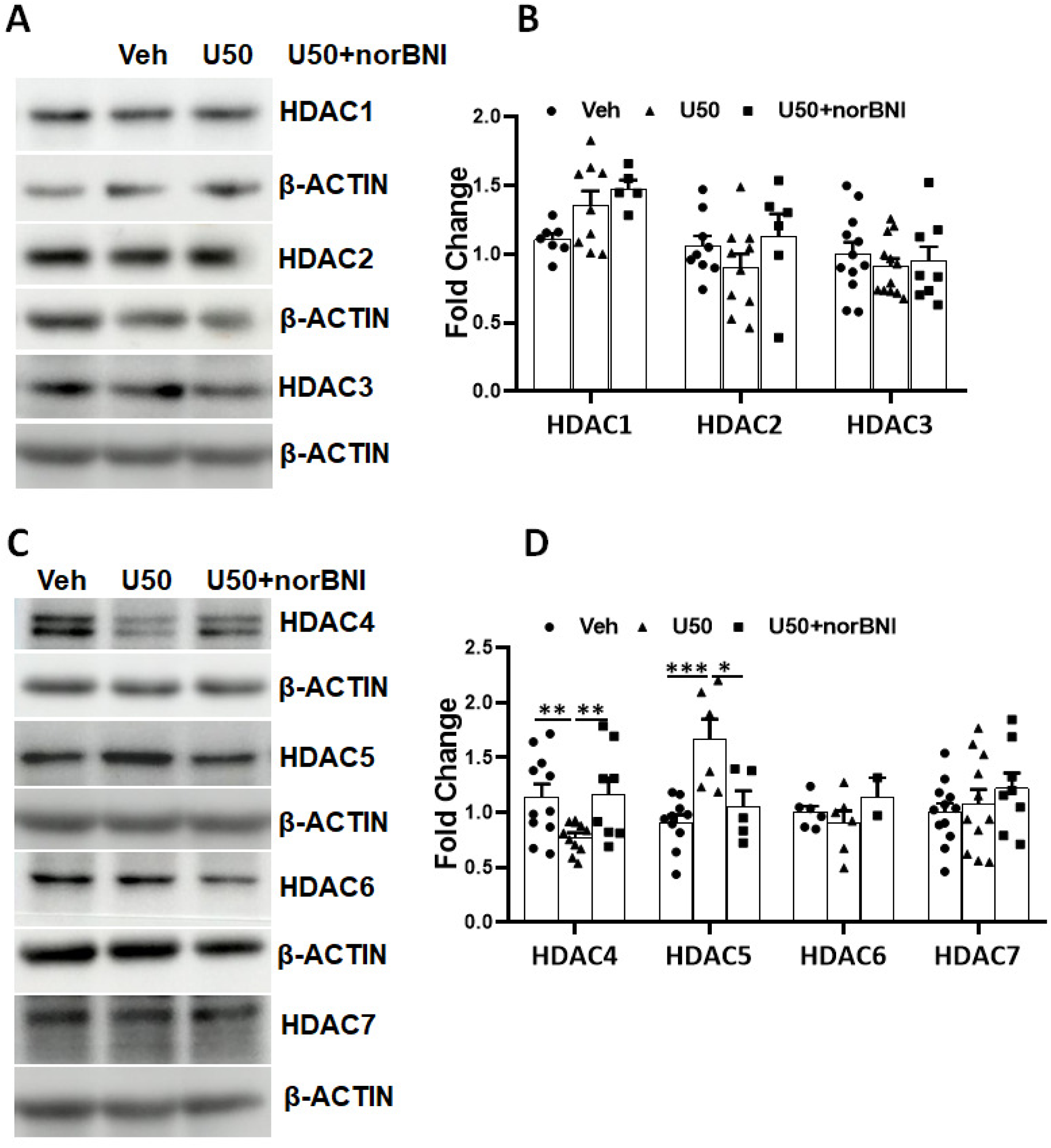
Effect of chronic KOR activation on histone deacetylases (HDACs) expression in the PFC. (**A**) No significant changes were observed in the expression of class I HDACs (HDAC1, HDAC2, and HDAC3) in the PFC after systemic treatment with U50488 (U50; 5 mg/kg; *i.p*.; 21 days). (**B**) Bar graph showing the quantification of the immunoblots presented in panel (A), n= 6-8 mice/group. (**C**) Significant decreased levels of HDAC4, and increased levels of HDAC5 expression were observed in the PFC of U50488-treated mice as compared to the Vehicle (Veh)-treated mice. Treatment with KOR antagonist, norBNI (10 mg/kg; *i.p.*; single injection on day-1 of the U50488 treatment) significantly blocked the effect of U50488 treatment on HDAC4 and HDAC5 levels in the PFC. No changes in the levels of HDAC6 and HDAC7 were observed in the PFC after chronic treatment with U50488. **(D**) Bar graph depicting the quantification of immunoblots presented in panel (C). Data are presented as mean ± SEM (^#^*p*<0.05, **^, ##^*p*<0.01, and \*\*\**p*<0.001 by Student’s t-test; For HDAC4: Veh vs U50 *t*= 3.045, U50 vs U50+norBNI, *t*= 2.911, n= 8-10; For HDAC5: Veh vs U50 *t*= 3.91264, U50 vs U50+norBNI, *t*= 2.527, n= 5-7).

**Figure 4.**
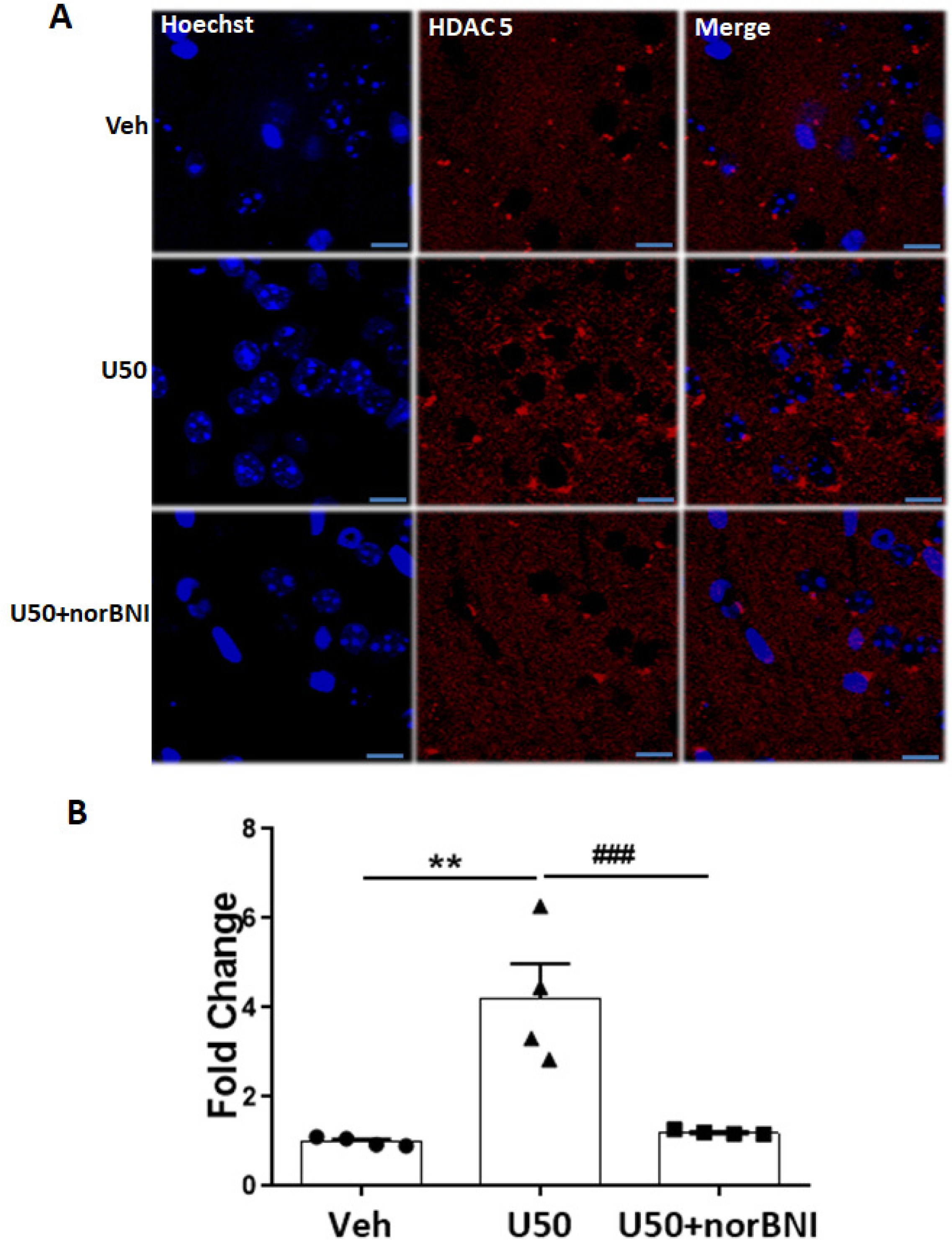
Sustained activation of KOR increased the expression of HDAC5 in anterior cingulate cortex. (**A**) Representative immunohistochemistry images showing the expression of HDAC5 (Red) and nucleus stained with Hoechst 33258 (Blue) in the anterior cingulate cortex (ACC) after treatment with U50488 (U50, 5 mg/kg; i.p.; 21 days), Scale bar= 20 µm. (**B**) Bar graph showing the quantification of HDAC5 expression presented in panel (A). (\*\**p*= 0.0009, *F*_(2,9)_= 16.64, \*\**p*<0.01, ^###^*p*<0.001 by One way ANOVA followed by Newman-Keul’s multiple comparisons test, n= 4/group).

### 4. HDAC5 regulates the expression of *Bdnf II* and *IV* transcripts

To further understand the relationship between HDAC5 and the expression of various *Bdnf* transcripts in the PFC, we performed a ChIP assay with an anti-HDAC5 antibody and determined the enrichment of HDAC5 at various *Bdnf* promoters. We observed that HDAC5 binding was significantly increased in *Bdnf II* and *IV* promoters in chronic U50488 treated mice as compared to vehicle-treated mice and treatment with norBNI blocked U50488-induced effects. Further, we did not observe any effects of U50488 treatment on HDAC5 binding at *Bdnf* promoters *I, III*, and *V* as compared to the vehicle treatment (**Figure 5A**). To confirm the contribution of HDAC5 enrichment on selective usage of *Bdnf II* and *IV* transcripts for BDNF synthesis, we evaluated the effect of a selective HDAC5 inhibitor, LMK235 (1 µM, 72 hours), on U50488-induced reduction in various *Bdnf* transcripts in mouse primary cortical neurons. As expected, LMK-235 completely blocked the inhibitory effects of U50488 treatment on *Bdnf II, IV* and *CDS* transcripts, albeit increased more than vehicle (**Figure 5B**).

**Figure 5.**
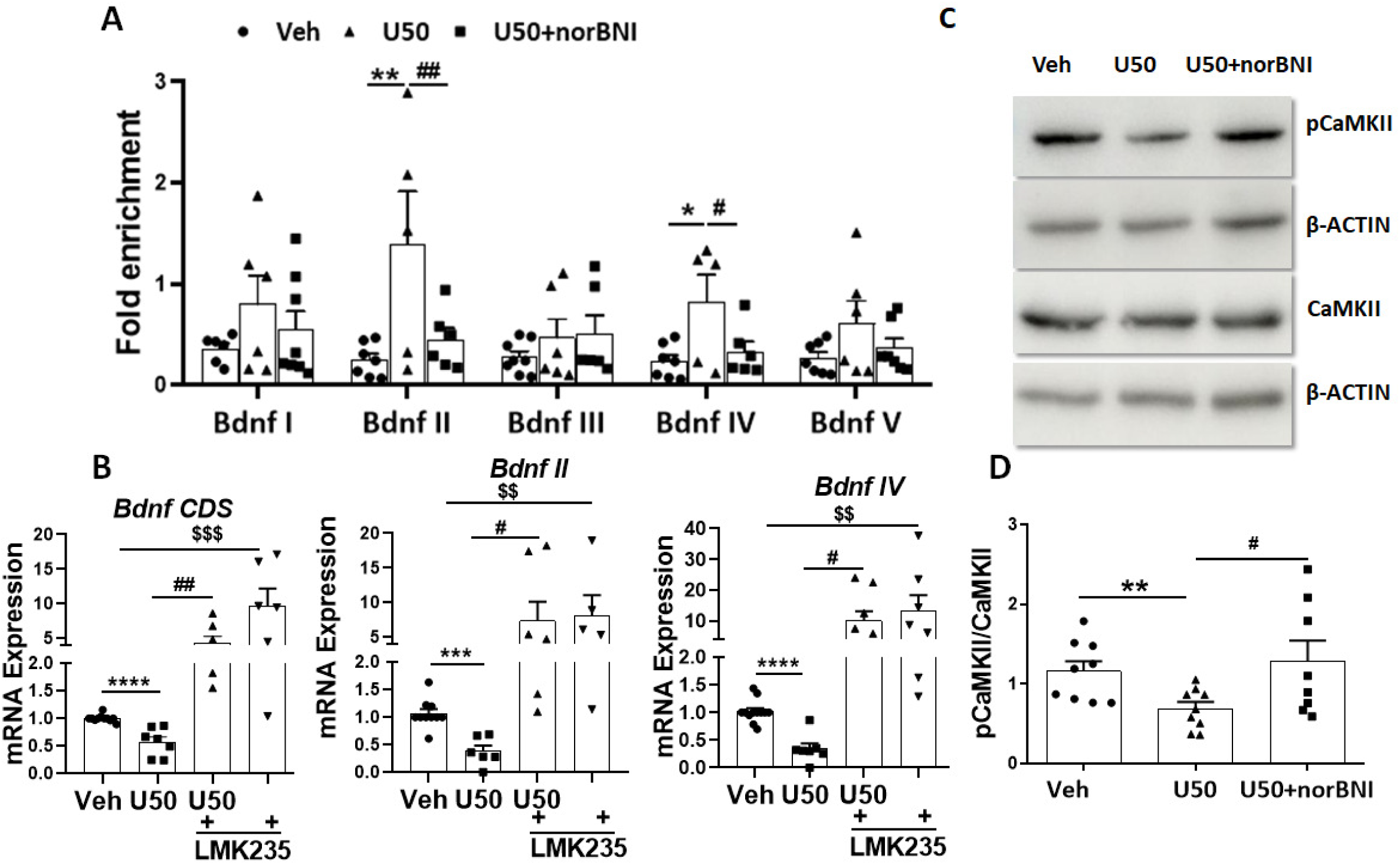
HDAC5 differentially regulates *Bdnf* transcripts in the PFC due to chronic KOR activation. (**A**) Chromatin immunoprecipitation (ChIP) with anti-HDAC5 shows a significant enrichment of HDAC5 binding at *Bdnf II* and *IV* promoters in the PFC after chronic administration of the KOR agonist, U50488 (U50; 5 mg/kg; *i.p*.; 21 days) as compared to the vehicle (Veh) treatment. Treatment with U50488 did not affect the levels of HDAC5 enrichment at *Bdnf I, II and V* promoters. Pre-treatment with a KOR selective antagonist, norBNI (10 mg/kg; *i.p.*; single injection on day-1 of the U50488 treatment), significantly blocked the effecst of U50488 treatment on HDAC5 enrichment at *Bdnf* promoters *II* and *IV*. (*Bdnf II* (n= 5-7): *p*=0.0194, *F*_(2,15)_= 5.814; *Bdnf IV* (n= 6-7): *p*= 0.0447, *F*_(2,16)_= 3.799; *^,#^*p*<0.05, **^,##^*p*<0.01 by One way ANOVA followed by Newman-Keul’s multiple comparisons test). Data are presented as mean ± SEM of fold change over input. (**B**) Pre-treatment of HDAC5 inhibitor, LMK235 (1 µM, 72 hours), in the primary cortical neurons blocked the U50488 (100 nM; for 72 hours)-induced decreases in the mRNA levels of *Bdnf CDS, II and IV* (*^, #, *$*^*p*<0.05, **^,##, *$$*^*p*<0.01, and ***^,###,*$$$*^*p*<0.001, ****^,####,*$$$$*^*p*<0.0001 by Student’s t-test, *Bdnf CDS:* Veh vs U50 *t*= 5.408; U50 vs U50+LMK235 *t*= 3.698; Veh vs LMK235 *t*= 4.687; *Bdnf II:* Veh vs U50, *t*= 5.139; U50 vs U50+LMK235, *t*= 2.308: Veh vs LMK235 *t*= 3.444; *Bdnf IV:* Veh vs U50, *t*= 6.373: U50 vs U50+LMK235, *t*= 2.866: Veh vs LMK235 t= 3.277; n= 7-9/group. (**C**) Mice treated with U50488 (U50, 5 mg/kg, 21 days, *i.p.*) exhibited significant suppression of pCaMKIIα (Thr286) levels in PFC as compared to the vehicle group, and this effect of U50488 was significantly reversed by KOR antagonist norBNI. (**D**) The bar graph showing the quantification of the pCaMKIIα expression normalized to total CaMKIIα levels of the immunoblots presented in panel (D) (\**p<*0.05, \*\**p<*0.01 by Student’s t-test; pCaMKIIα: Veh vs U50 t= 3.075; U50 vs U50+norBNI t= 2.381 n= 8-9 mice/group). (**E**) Chronic administration of U50488 significantly decreased the levels of myocyte enhancer factor 2C (MEF2C) in the PFC as compared to the vehicle treatment. Treatment with a selective KOR antagonist, norBNI, significantly blocked the effect of U50488 on the MEF2C levels in the PFC. (**F**) Bar graph depicting the quantification of immunoblots presented in panel € (\*\*\**p*<0.001, ^#^*p*<0.05 by Student’s t-test; Veh vs U50 *t*= 4.574, U50 vs U50+norBNI *t*= 2.637), n= 5-7 mice/group.

Both HDAC4 and HDAC5 interact with myocyte enhancer factor-2 (MEF2) and prevent the expression of its target genes, including *Bdnf* [24, 25]. It is also known that CaMKII activation dissociates MEF2C from HDAC4/5 and restores MEF2C transcriptional activity [26]. Further, CaMKII-dependent phosphorylation of nuclear HDAC5 leads to its translocation from the nucleus to cytoplasm and hence modulates its transcription repressive activity [27]. In light of the above mentioned facts, we determined the expression of pCaMKII (Thr286) in the PFC of mice treated with either vehicle, U50488, or U50488 + norBNI. We found significantly decreased expression of pCaMKII in U50488-treated mice as compared to the vehicle-treated mice (**Figure 5C, D**). Further, treatment with a selective KOR antagonist, norBNI completely blocked the effect of U50488 on the levels of pCaMKII in the PFC (**Figure 5C, D**). These findings indicate that increased HDAC5 expression after chronic treatment with selective KOR agonist,U50488, could be due to insufficient level pCaMKII not being able to phosphorylate HDAC5 and not promoting its export from nucleus and / or degradation as reported earlier [28]. These results suggest that sustained KOR activation reduced CAMKII phosphorylation in the PFC, which in turn reduces nuclear export of HDAC5 and its degradation. Considering the ability of MEF2C to regulate HDAC-induced regulation of *Bdnf* transcription and to modulate neuronal depolarization induced expression of *Bdnf exon IV* [29], we also determined the effect of chronic U50488 treatment on the expression of MEF2C in the PFC. As per our hypothesis, we observed significant inhibition of MEF2C expression in the PFC of U50488-treated mice as compared to the vehicle-treated mice, and this effect of U50488 was blocked by KOR antagonist, norBNI, suggesting KOR-specific effect (**Figure 5E, F**). These results clearly suggest that sustained KOR signaling in the PFC lead to the suppression of pCaMKIIα levels, due to which HDAC5 is retained in the nucleus to deacetylate Histone H3 at Lysine 9 residue, and consequently inhibits MEF2C mediated transcription of *Bdnf I* and *IV* exons.

### PFC HDAC5 is essential for KOR mediated depression

Based on our findings that KOR activation significantly elevated HDAC5 expression and suppressed *Bdnf II* and *IV* transcripts in the PFC, we sought to determine if HDAC5 is essential for KOR-induced downregulation of the *Bdnf* expression and induction of depression-like behaviors. We knocked down HDAC5 in the PFC by employing stereotaxic delivery of adeno associated viral particles containing *HDAC5*-shRNA (AAV-*HDAC5*-shRNA; **Figure 6A**) in adult C57BL/6 mice (**Figure S9**). We observed significantly decrease in the levels of *Hdac5* mRNA in the PFC of mice expressing *HDAC5*-shRNA (**Figure 6B**). Further, knock-down of HDAC5 in the PFC completely blocked U50488-mediated suppression of *Bdnf CDS, Bdnf II,* and *Bdnf IV transcripts* in the PFC of mice expressing *HDAC5*-shRNA as compared to the scrambeled-shRNA expressing mice (**Figure 6 B-E**). We next evaluated the contribution of HDAC5 in mediating KOR-induced behavioral deficits relevant to depressive behaviors using forced swim test, tail suspension test and 3-chamber social behavior test. We observed that HDAC5 knockdown completely blocked U50488-induced increases in immobility time in forced swim test (**Figure 6F**) and the tail suspension test (**Figure 6G**). HDAC5 knockdown also rescued chronic KOR activation-induced deficits in sociability (**Figure 6H**). These results further confirm that sustained KOR activation-induced increases in the levels of HDAC5 in the PFC are critical for KOR-induced decrease in *Bdnf* transcription and induction of depression-like behaviors in mice.

**Figure 6.**
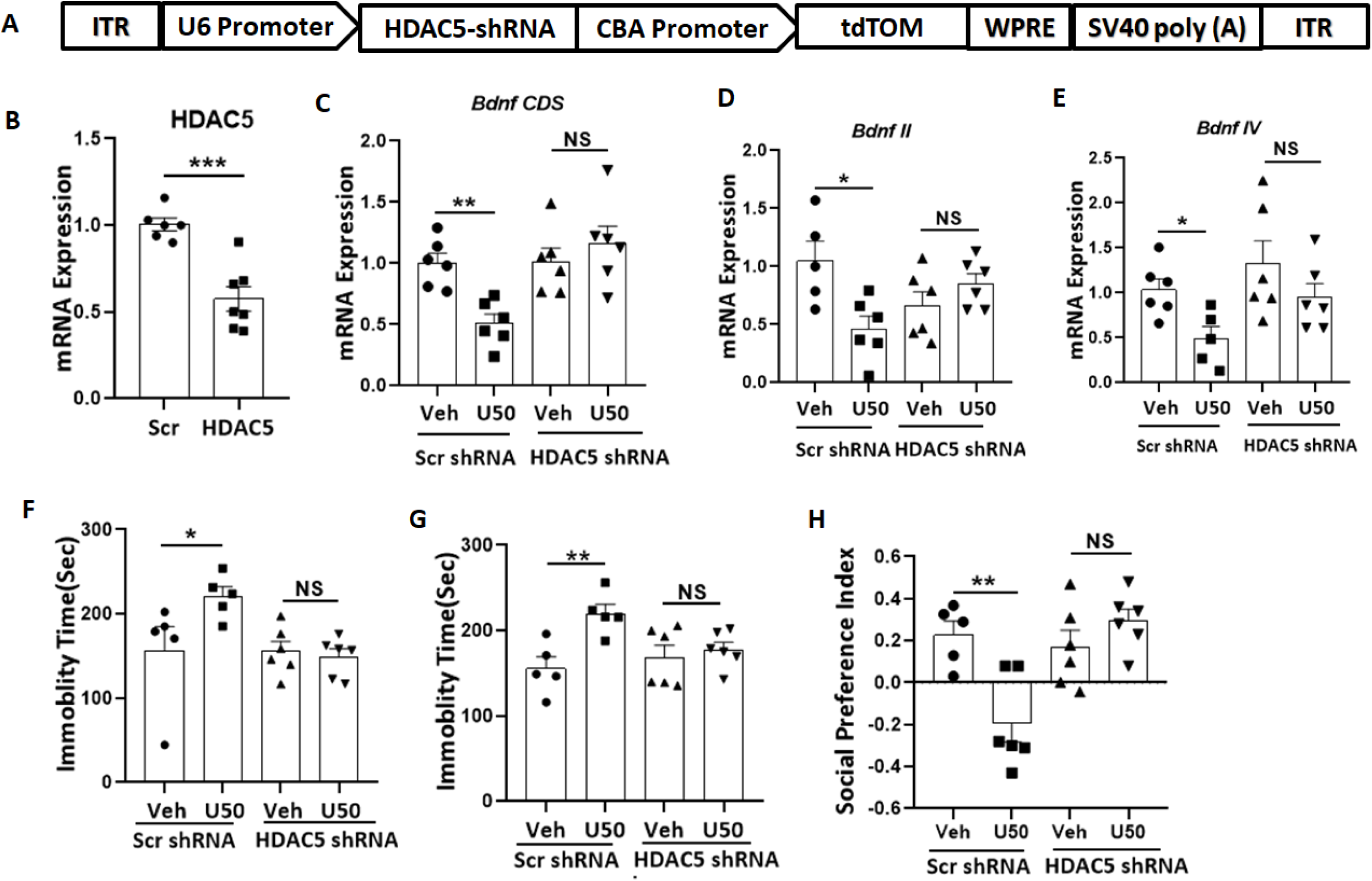
HDAC5 knockdown in the PFC blocked the effects of the KOR agonist, U50488, on various *Bdnf* transcripts and depression-like behaviour. (**A**) Schematic diagram showing the arrangement HDAC5-shRNA or scrambled-shRNA (Scr shRNA) in adeno-associated viral vector. (**B**) Bar graph showing extent of HDAC5-shRNA-mediated knockdown of *Hdac5* mRNA in the PFC (***p<0.001, Scr-shRNA vs HDAC5-shRNA, Student’s t-test; *t*=5.189, n= 6-7 mice/group). (**C, D, E**) Knockdown of HDAC5 in the PFC significantly blocked chronic U50488 treatment (U50; 5 mg/kg *i.p.*; 21 days)-induced reductions in the levels of *Bdnf CDS, Bdnf II* and *Bdnf IV* (*, *p*<0.05, and \*\**p*<0.01, by Student’s t-test *(Scrembled Veh vs Scrembled U50,* t=4.49); *Bdnf II (Scrembled Veh vs Scrembled U50*, *t*=3.042) and *Bdnf IV (Scrembled Veh vs Scrembled U50, t*=3.001) transcripts in the PFC, N=6 mice/group). (**F**) Chronic administration of U50488 increased the immobility time measured in the forced swim test (FST) in the mice expressing scrambled-shRNA in the (* *p*<0.05, by Student’s t-test (Veh vs U50, *t*=2.978). The effects of U50488 on the immobility time in the FST were absent in the mice expressing HDAC5 in the PFC. (**G**) Chronic administration of U50488 increased the immobility time in the tail suspension test (TST) in the mice expressing scrambled-shRNA in the PFC (** *p*<0.01, by Student’s t-test (Veh vs U50, *t*=3.732). The effects of U50488 on the immobility time in the TST were absent in the mice expressing HDAC5 in the PFC. (**H**) In a 3-chamber social behaviour test, chronic administration of U50488 decreased the social preference index in the mice expressing scrambled-shRNA in the PFC (**, *p*<0.01, by Student’s t-test (Veh vs U50, *t*=3.698). The effects of U50488 on the social preference index were absent in the mice expressing HDAC5 in the PFC.

## Discussion

The KOR is widely expressed in the CNS and plays an important role in regulating mood, pain, and addiction [30]. The inhibitory effect of increased KOR/Dyn signaling on monoaminergic neurotransmission in various models of psychiatric disorders, including major depression, is widely known [31, 32]. Importantly, several clinical studies with KOR antagonist in depressed patients [15, 16, 33], strongly advocate the KOR as tractable target for TRD. Although increased BDNF expression in the PFC and the hippocampus is considered as a vital transducer of the antidepressant response [34], the mechanistic relationship between KOR signaling and BDNF expression is not completely understood. In this study, we demonstrate that sustained activation of KOR decreases *Bdnf* mRNA (protein-coding transcript *exon IXa*) expression via suppression of *Bdnf II* and *Bdnf IV* transcripts that act as alternative promoters. Importantly, we also observed a significant decrease in the levels of active chromatin marker H3K9ac and increase in the levels of HCAC5 in the PFC of mice that were chronically administered with KOR agonist, U50488. Our mechanistic studies using ChIP analysis performed with anti-HDAC5 and anti-H3K9ac antibodies showed decreased binding of H3K9ac and significant enrichment of HDAC5, at BDNF *II* and *IV* promoters in the PFC of mice chronically treated with U50488. Our study also revealed that chronic KOR activation with U50488 decreases the PFC levels of pCaMKIIα, which is known to phosphorylate HDAC5 leading to its nuclear export and degradation, a mechanism that ultimately facilitates BDNF transcription. We also observed decreased expression of transcription factor MEF2C that has been shown to regulate expression of *Bdnf IV* transcript. Optimal BDNF expression is essential for a wide array of neurological functions, including neuronal survival and differentiation, synaptic neurotransmission, and synaptic consolidations [35, 36], which is spatiotemporally regulated by transcription of each 5’-exon controlled by a separate promoter [37] and several transcription factors [38]. Activity-dependent changes in mouse *Bdnf-I, IV*, or *IX* transcripts in the PFC and in the hippocampus have been reported [39–41]. The importance of regulation of *Bdnf* expression via changes in non-coding *Bdnf* transcripts in major depression is further highlighted by a recent study [42] that demonstrated that *Bdnf* L-3’-UTR mediates dendritic shrinkage and depressive and anxiety-like behaviors in humans and mice. However, there is no study on the levels of other major non-coding *Bdnf* transcripts in treatment-resistant depression-like condition observed after chronic treatment with U50488. In this context, our findings showing significant changes in *Bdnf II* and *IV* transcripts in the PFC of U50488-treated mice shed light on the complex regulation of BDNF expression in depressive disorders. *Bdnf IV* plays an essential role in the regulation of dendritic integrity [37] and GABAergic neurotransmission in the PFC [43] both of which are altered in depressed patients and rodents modeling depression-like symptioms [44, 45]. Further, hypomethylation of *Bdnf IV* is also associated with non-remission by antidepressants [46]. In light of these studies, our observation of KOR-induced repression of *Bdnf II* and *IV* transcription could provide insights into mechanisms involved in maladaptive synaptic plasticity and neurotransmission underlying treatment-resistant depression.

Changes in histone acetylation have been widely implicated in the pathogenesis of several neurological disorders, including major depression [47]. Earlier class II HDAC, HDAC5 has been shown to epigenetically regulate the behavioural adaptations in the nucleus accumbens in response to chronic stress [48] and to block the antidepressant-like effects of imipramine in the chronic social defeat stress model of depression [12, 49]. Furthermore, ketamine-induced fast antidepressant response has been shown to be mediated through CaMKIIα-induced HDAC5 phosphorylation and its subsequent nuclear export, leading to the activation of several target genes (e.g., MEF2, EGR1, etc.) in the hippocampus [50]. In light of these studies, our finding that chronic activation of KOR increases the levels of HDAC5 in the PFC, and that knockdown of HDAC5 in the PFC completely blocks the effects of KOR activation on *Bdnf II* and *IV* transcripts uncovers a new facet of KOR signaling and its role in depressive disorders. The contrasting effects of KOR activation on HDAC4 and HDAC5 could be attributed to varying cell type (GABAergic vs Glutamatergic neurons) and activity status as even spontaneous neuronal activity has been shown to decrease the HDAC4, but not of HDAC5 [51]. We did not observe any effect of KOR stimulation on the levels of the HDAC5 in the hippocampus (data not presented), which is in contrast to chronic social defeat model of depression [12, 49]. These discrepancies may stem due to KOR being expressed in different neuronal populations in the PFC and the hippocampus [52]. Nevertheless, KOR induced HDAC5 and decreased H3K9ac expression in the PFC appears to be pivotal for downstream events such as decreased expression of MEF2C and BDNF levels. Since HDAC5 shuttling between the nucleus and the cytoplasm depends on its phosphorylation by CaMKII [49], our results revealing decreased levels of pCaMKII in the PFC, increased HDAC5 binding at *Bdnf II* and *Bdnf IV* promoters, and decreased enrichment of H3K9ac at *Bdnf II* and *IV*, are consistent with the prevailing notion of epigenetic silencing of *Bdnf* in major depression [53, 54].

According to the Diagnostic Statistical Manual (DSM-V) major depressive disorder (MDD) is diagnosed as a single entity, however, there is a huge array of symptoms that could also meet the DSM-V criteria, clearly depicting the heterogeneity of etiology and pathophysiology of MDD [55]. Given that KOR modulates all three major neurotransmitters, dopamine, serotonin, and norepinephrine, in the brain [56], even subtle over-activation could lead to severe mood alteration via engaging multiple neurocircuits and signaling pathways, which could be resistant to pharmacotherapies selectively targeting one or other pathways. Furthermore, our data of KOR being a selective modulator of HDAC5 activity in the PFC further supports KOR as a druggable target to selectively modulate epigenetic signaling in depressive disorders, which is otherwise impossible to target due to a lack of tissue and cell selectivity. In conclusion, our data of KOR induced upregulation of HDAC5 in PFC further supports the hypothesis of perturbation of multiple pathways in depression, and hence looking beyond monoaminergic targets, such as KOR for the treatment of MDD could be more fruitful.

## Supporting information

supplementary Information.docx

## Acknowledgments

The authors thank Mrs. Deepmala Umrao and Mr. Devanshu Kaushik for their editorial help. We also acknowledge Intravital Imaging Facility Olympus BX61-FV1200-MPE. This manuscript bears CSIR-CDRI communication number 99/2023/PNY.

## Author Contributions

Anubhav Yadav-Manuscript writing, Contribution to figure 1(A& B), 2(E), 3(A, B, C&D), 4(A, & B), 5(A, B,C, and D), 6(A,B,C,D,E,F,G and H), S1(A,&B), S2(A,B&C), S1(A,&B), S5 and S6(A&B), Shalini Dogra-Manuscript writing, Contribution to figure 1(A& B), 2(A, B, C and D), S2(A, B&C), S3(A, B, C, D, E, F, G&H), S4(A, &B), and S5, Poonam Kumarif-Analysis of ChIP data, and dissection of mice brain, Boda Arun Kumar-Optimizations of ChIP assay, Ajeet Kumar-Determining the effect of KOR agonist in a rodent model of chronic unpredictable stress, Manish Kumar Dash-Brain perfusion and immunohistochemistry, Prem Narayan Yadav-Design of experiments, data interpretation and manuscript writing

