## supplementary Information.docx for "Kappa opioid receptor activation induces epigenetic silencing of brain-derived neurotropic factor via HDAC5 in depression"

**Table S1.** List of primers used for RT-PCR or ChIP-PCR.

| **S.N.** | **Primer name** | **Sequence (5' - 3')** |
|---|---|---|
| **Primers used for CHIP-PCR** | | |
| 1 | *Bdnf I*-F | TTGATCATCACTCACGACCACG |
| 2 | *BdnfI*-R | CAGCCTCTCTGAGCCAGTTACG |
| 3 | *Bdnf II*-F | CCGTCTTGTATTCCATCCTTTG |
| 4 | *BdnfII*-R | CCCAACTCCACCACTATCCTC |
| 5 | *Bdnf III*-F | GTGAGAACCTGGGGCAAATC |
| 6 | *Bdnf III*-R | AAAGAACGGAAAAGAGGGAG |
| 7 | *Bdnf IV*-F | CTTCTGTGTGCGTGAATTTGCT |
| 8 | *BdnfIV*-R | AGTCCACGAGAGGGCTCCA |
| 9 | *Bdnf V*-F | TGAGACTCACACTCGCTTCCTC |
| 10 | *Bdnf V*-R | GCACTGGCTTCTCTCCATTTC |
| **Primers used for mRNA analysis** | | |
| 11 | β-Actin F | TGTTACCAACTGGGACGACA |
| 12 | β-Actin R | CTGGGTCATCTTTTCACGGT |
| 13 | *Bdnf I*-F | CCTGCATCTGTTGGGGAGAC |
| 14 | *Bdnf* I and IV-R | GCCTTGTCCGTGGACGTTTA |
| 15 | *Bdnf* II-N-F | TTGGCTTCCTAGCGGTGTAGG |
| 16 | *Bdnf* II and V-R | AGGATGGTCATCACTCTTCTCACCT |
| 17 | *Bdnf* V-F | CTTGGGGCAGACGAGAAAGC |
| 18 | *Bdnf* IV-R | CAGAGCAGCTGCCTTGATGTT |
| 19 | *Bdnf III*-F | GTGAGAACCTGGGGCAAATC |
| 20 | *Bdnf III*-R | AAAGAACGGAAAAGAGGGAG |
| 21 | *Bdnf* CDS-F | AGGTGAGAAGAGTGATGACCATCC |
| 22 | *Bdnf* CDS-R | AGCCAGTGATGTCGTCGTCAG |
| 23 | *GAPDH- F* | GAAGGTCGGTGTGAACGGAT |
| 24 | *GAPDH- R* | ACTGTGCCGTTGAATTTGCC |
| 25 | *Hdac5-F* | TGTCACCGCCAGATGTTTTG |
| 26 | *Hdac5-R* | TGAGACGAGCCGAGACACAG |

**Table S2.** List of antibodies and its dilution used in various experiments

**Table S2.** List of antibodies and their dilution used in various assays.

| Antibody Name | Dilutions | Source | Catalog No. | Application |
|---|---|---|---|---|
| Anti-H3K9ac | 1:1000 | Abcam | ab10812 | WB/ChIP/IHC |
| Anti-H3K4me3 | 1:1000 | Abcam | ab12209 | WB |
| Anti-H3K9me3 | 1:1000 | Abcam | ab8898 | WB |
| Anti-H3K27me3 | 1:1000 | Abcam | ab6002 | WB/ ChIP |
| Anti-Total Histone | 1:1000 | Abcam | ab1791 | WB |
| Anti-HDAC1 | 1:1000 | Cell Signaling | #3458 | WB |
| Anti-HDAC2 | 1:1000 | Cell Signaling | #5715 | WB |
| Anti-HDAC3 | 1:1000 | Cell Signaling | #8505 | WB |
| Anti-HDAC4 | 1:1000 | Cell Signaling | #1516 | WB |
| Anti-HDAC5 | 1:1000 | Cell Signaling | #2045 | WB |
| Anti-HDAC5 | 1:50 | Proteintech | 16166-1-AP | ChIP/IHC |
| Anti-HDAC6 | 1:1000 | Cell Signaling | #7612 | WB |
| Anti-HDAC7 | 1:1000 | Cell Signaling | #3341 | WB |
| Anti-HDAC8 | 1:1000 | Cell Signaling | #6604 | WB |
| Anti-SIRT1 | 1:1000 | Cell Signaling | #9475 | WB |
| Anti-SIRT2 | 1:1000 | Cell Signaling | #1265 | WB |
| Anti-SIRT3 | 1:1000 | Cell Signaling | #5490 | WB |
| Anti-SIRT4 | 1:1000 | Cell Signaling | #6978 | WB |
| Anti-SIRT5 | 1:1000 | Cell Signaling | #8782 | WB |
| Anti-SIRT6 | 1:1000 | Cell Signaling | #1248 | WB |
| Anti-SIRT7 | 1:1000 | Cell Signaling | #5360 | WB |
| Anti-CaMKIIα | 1:1000/1:50 | Thermo-fisher Scientific | MA1-048 | WB/IHC |
| Anti -pCaMKIIα | 1:1000 | Thermo-fisher Scientific | MA1-047 | WB |

**Supplementary Figures:**

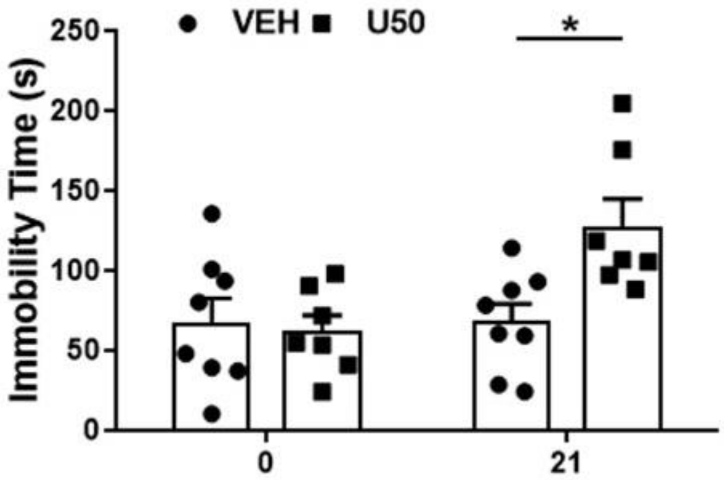


**Figure S1.** Chronic KOR activation induces depression in C57BL6/J mice. Chronic treatment of mice with a selective KOR agonist, U50488 (U50, 5 mg/mg/kg, *i.p.*) for 21 days led to significant increase in immobility time as compared to the vehicle (Veh) treatment as measured by forced swim test. $F_{(1,13)}$= 8.657; $p$= 0.0114 by Two Way ANOVA followed by Bonferroni's multiple comparisons test *$p$<0.05 (n= 7-8 mice/group). Data are presented as the mean ± SEM of % Veh.

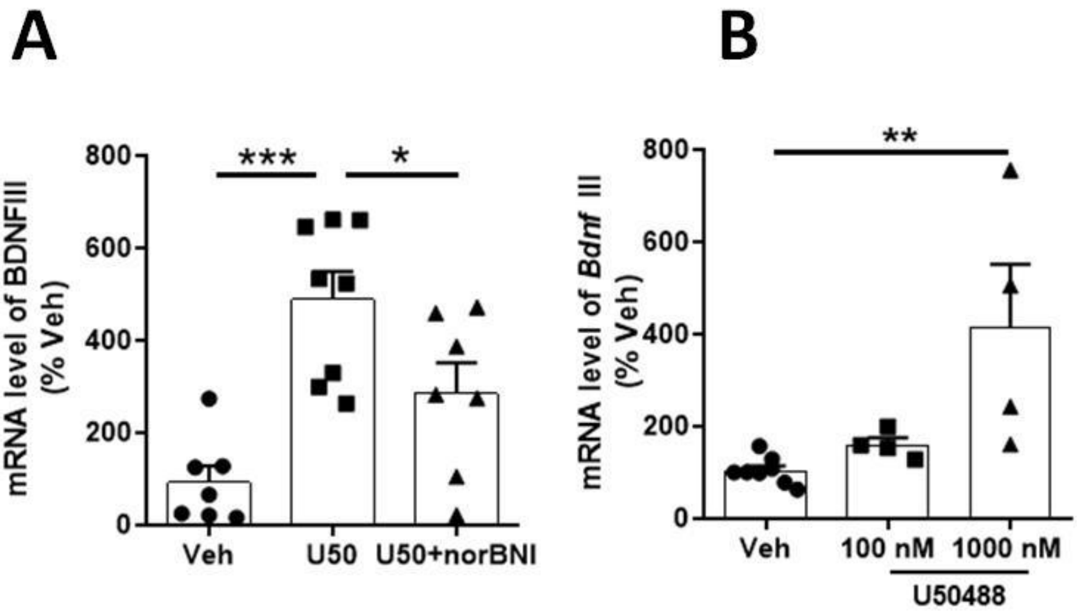


**Figure S2.** KOR activation leads to differential regulation of *Bdnf* transcripts. **(A)** Chronic treatment with U50488 (5 mg/kg; *i.p.*; 21 days) significantly increased the levels of *Bdnf* III transcript in mice's prefrontal cortex (PFC). A single administration of KOR antagonist, nor-binaltorphimine (norBNI, *i.p.*; 10 mg/kg), blocked the effects of U50488 treatment on the expression of *Bdnf III*. Data are presented as mean ± SEM of % Vehicle, $p<0.001$, $F_{(2,19)}= 13.03$, *$p<0.05$ (Veh vs U50), (U50 vs U50+norBNI) ***$p<0.001$ (Veh vs U50) (U50 vs U50+norBNI) by One way ANOVA followed by Newman-Keul's multiple comparisons test, n= 5-7 mice/group. (**B)** Representative bar graph showing increased expression of *Bdnf III*, transcript (normalized to GAPDH transcript) in 12 days *in vitro* (DIV12) primary cortical neurons treated with U50488 (100 nM and 1000 nM;72 hours). Data are presented as mean ± SEM of % Vehicle, $p<0.01$, $F_{(2,13)}= 7.629$, **$p<0.01$ (Veh vs 1000nM U50) by One way ANOVA followed by Newman-Keul's multiple comparisons test, n= 5-7/group.

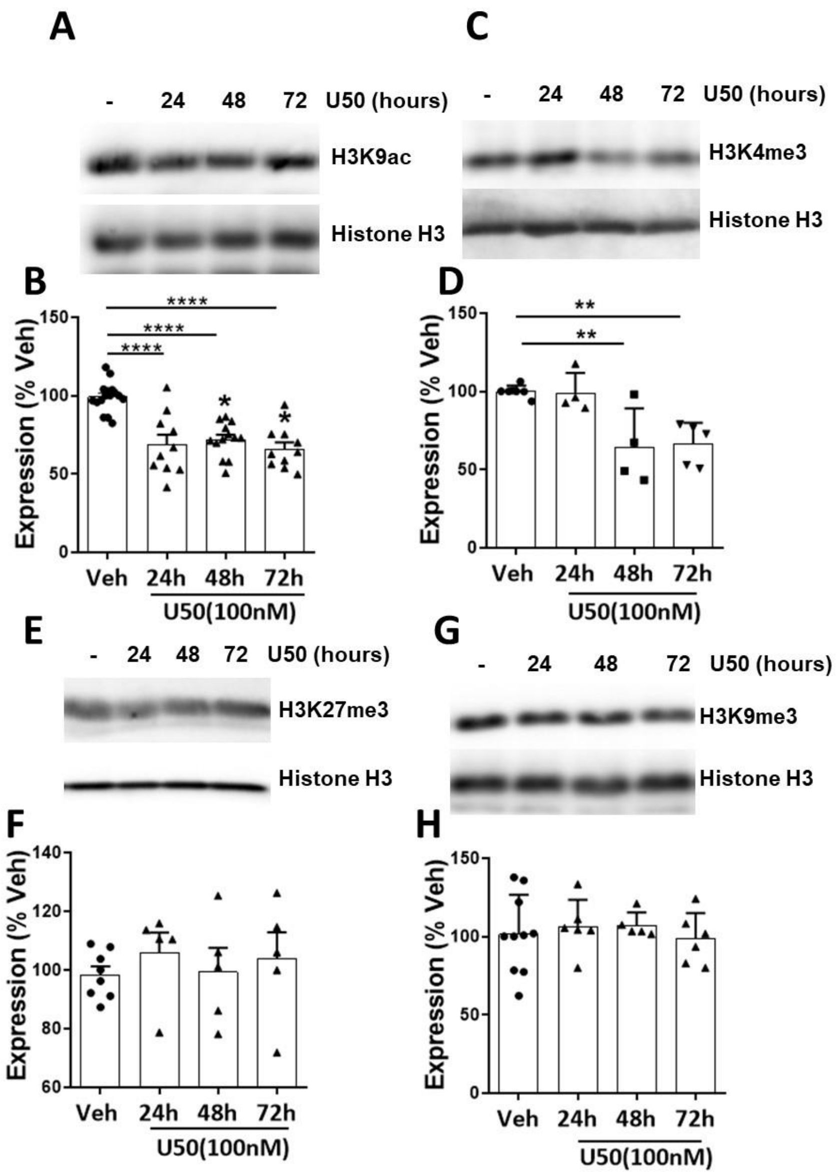


**Figure S3.** Immunoblots showing epigenetic modifications in the primary cortical neurons. (**A, B**) The time-dependent treatment (24, 48, and 72 hours) of U50588 (U50, 100 nM) significantly decreased the levels of acetylation at 9$^{th}$ lysine of the histone H3 (H3K9ac) in the primary cortical neurons (****$p$<0.0001, $F_{(3,45)}$= 19.01 by one way ANOVA followed by Newman-Keul's multiple comparisons test, n= 5-7/group test). (**C, D**) U50488 treatment for 48 and 72 hours decreased levels of tri-methylation at 4$^{th}$ lysine residue of the histone H3 (H3K4me3) in the primary cortical neurons (***$p$= 0.0006; $F_{(3,16)}$= 9.885 by one way ANOVA followed by Newman-Keul's multiple comparisons test, n= 5-7/group test). Data are presented as mean ± SEM of % Vehicle (Veh). (**E, F**) U50488 treatment did not affect the levels of tri-methylation at 27$^{th}$ lysine residue of the histone H3 (H3K27me3) in the primary cortical neurons. (**G, H**) No effect of U50488 treatment on the levels of tri-methylation at 9$^{th}$ lysine residue of the histone H3 (H3K9me3) in the primary cortical neurons.

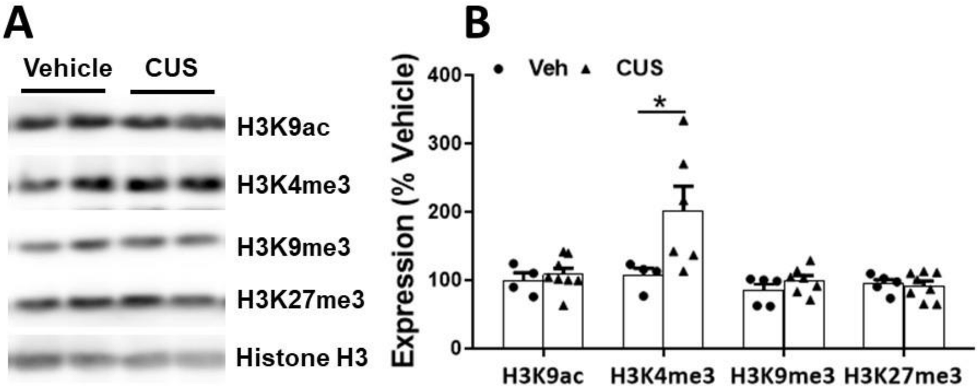


**Figure S4.** Exposure to chronic unpredictable stress (CUS) induces epigenetic changes in the prefrontal cortex (PFC). **(A)** Significant increase in the levels of tri-methylation at 4th lysine residue of the histone H3 (H3K4me3) was found after chronic unpredictable stress (CUS) treatment as compared to the vehicle treatment. No changes were observed in the levels of acetylation at 9th lysine residue of the histone H3 (H3K9ac), tri-methylation at 9th lysine residue of the histone H3 (H3K9me3), and tri-methylation at 27th lysine residue of the histone H3 (H3K27me3) in the PFC of CUS-exposed mice as compared to the vehicle treatment. (**B**) Bar graph showing the quantification of the western blots presented in panel (A) (*$p$<0.05, t= 2.518, by Student's t-test, n= 6-8 mice/group). Data are presented as mean ± SEM of % Vehicle.

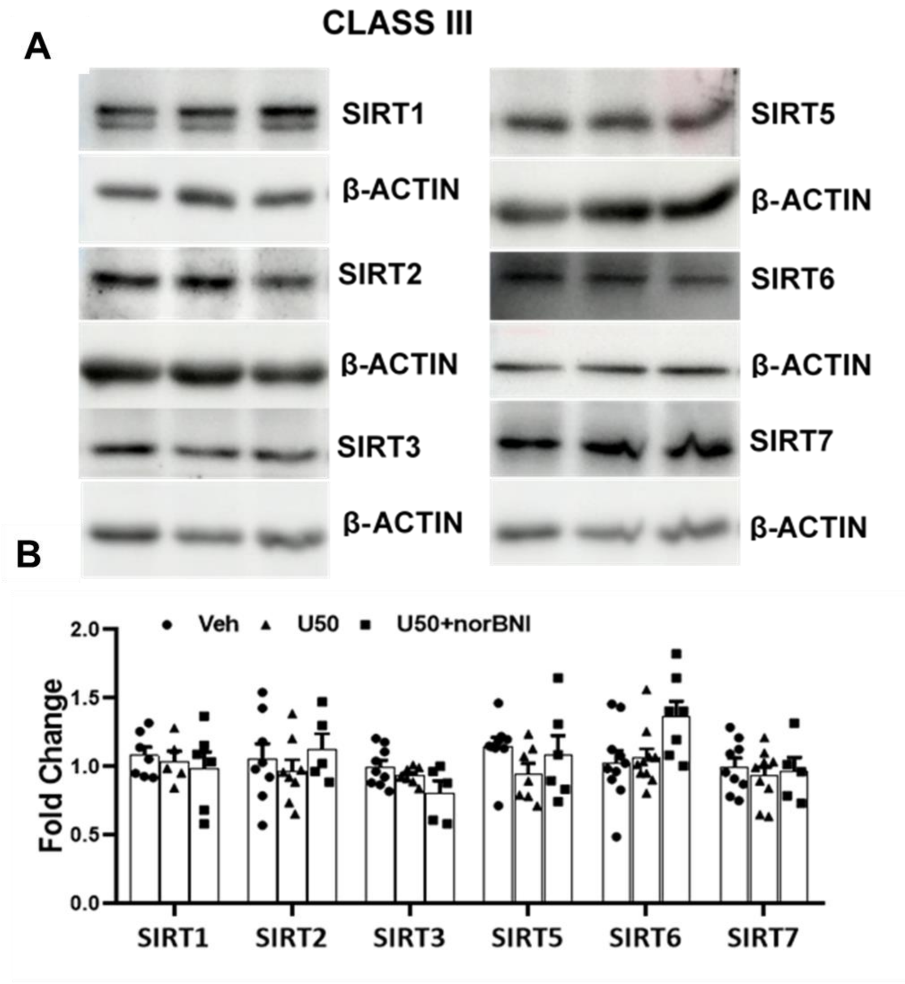


**Figure S5.** Effect of chronic U50488 treatment the levels of class-III family of HDACs. (**A**) Representative Immunoblots showing the effect of chronic U50488 (U50) and norbinaltorphimine (norBNI) treatment on the levels of class III HDACs. (**B**) Bar graph showing quantification of blots presented in panel (A). Data are presented as mean ± SEM of fold change over Vehicle treatment (n= 6-8 mice/group). Veh= Vehicle, SIRT1= Sirtuin 1, SIRT2= SIrtuin 2, SIRT3= Sirtuin 3, SIRT5= Sirtuin 5, SIRT6 =Sirtuin 6, SIRT7= Sirtuin 7

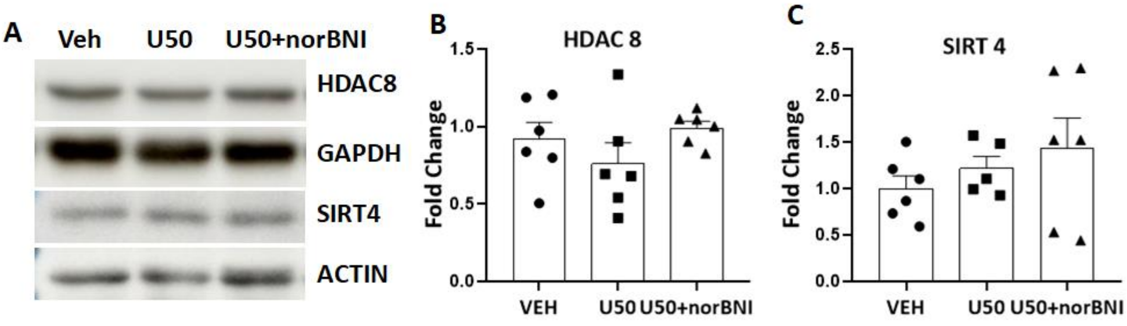


**Figure S6.** Effect of chronic KOR activation on HDAC8 and SIRT4 expression in the PFC. (**A**) No significant changes in the expression of HDAC8 and SIRT4 in the PFC were observed after chronic U50488 treatment (5 mg/kg; *i.p.*; 21 days). (**B,C**) Bar graph showing the quantification of the immunoblots presented in panel (A). Data are presented as mean ± SEM of fold change over Vehicle treatment (n=5-6 mice/group).

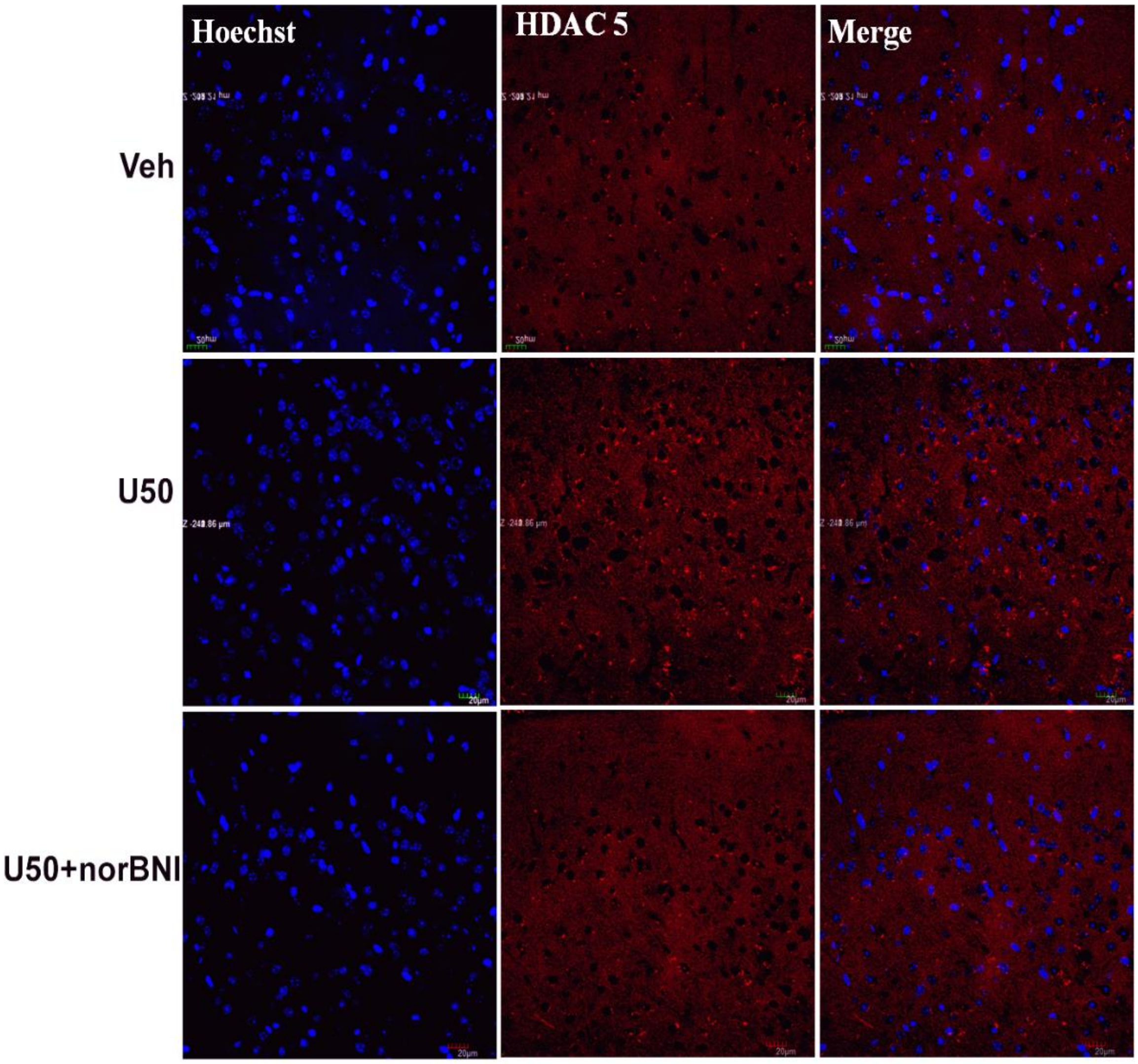


**Figure S7.** Representative confocal image of the brain section (Uncropped) depicting landmarks of anterior cingulate cortex showing the expression of HADC5 (red) and Hoechst 33258 (blue).

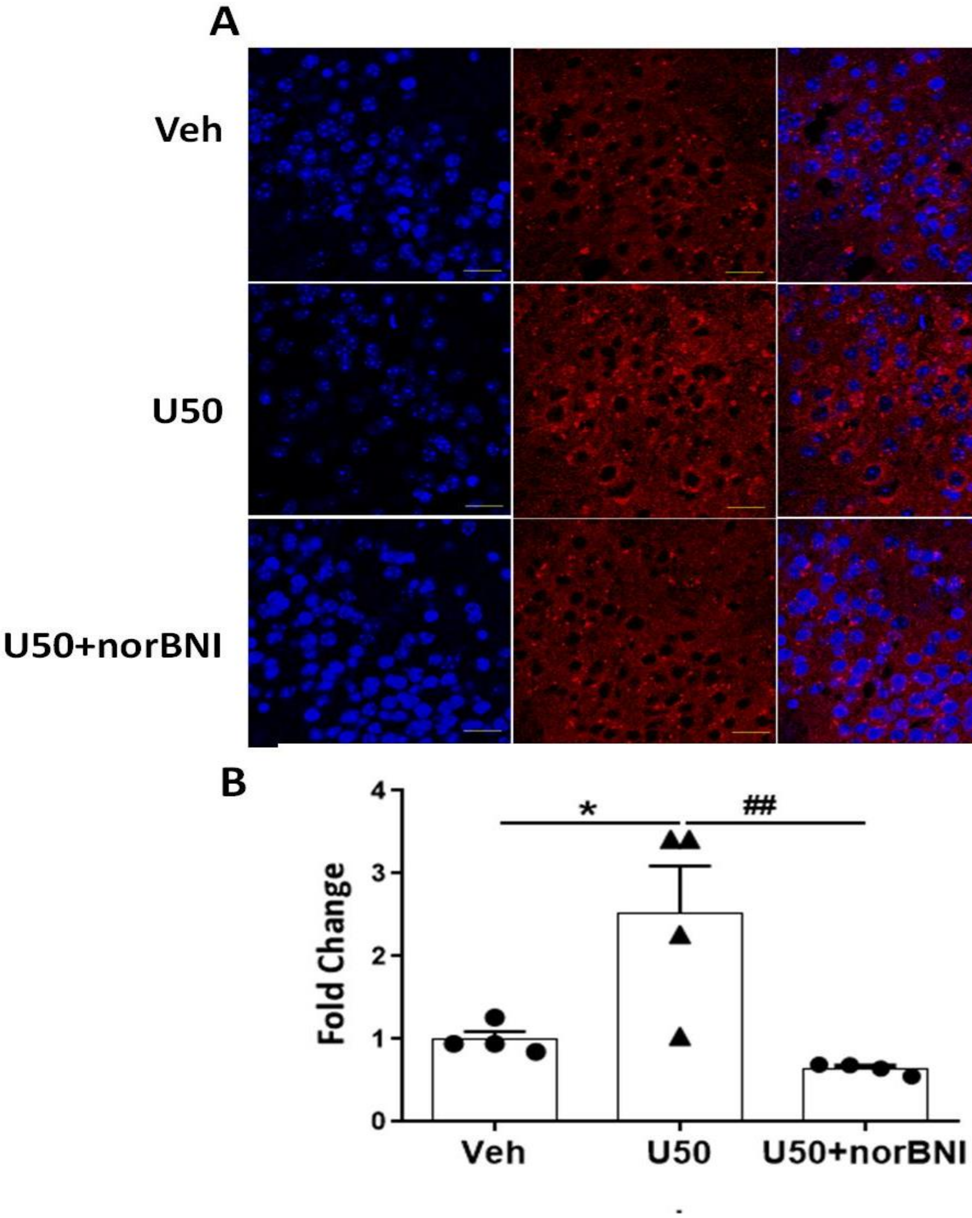


**Figure S8.** Sustained KOR activation attenuated HDAC5 expression in the piriform cortex. **(A)** Representative immunohistochemistry images showing the expression of HDAC5 (red) and nuclear staining with Hoechst 33258 (blue) in the piriform cortex of mice after various treatments. Scale bar= 20 µm (**B**) Bar graph showing the quantification of HDAC5 expression. Data are presented as mean ± SEM ($p$< 0.01, $F_{(2,9)}$= 9.018, *$p$<0.05, $^{\#\#}p$<0.01 by one way ANOVA followed by Newman-Keul's multiple comparisons test, n= 4/mice. Veh= Vehicle, U50= U50488, norBNI= norbinaltorphimine

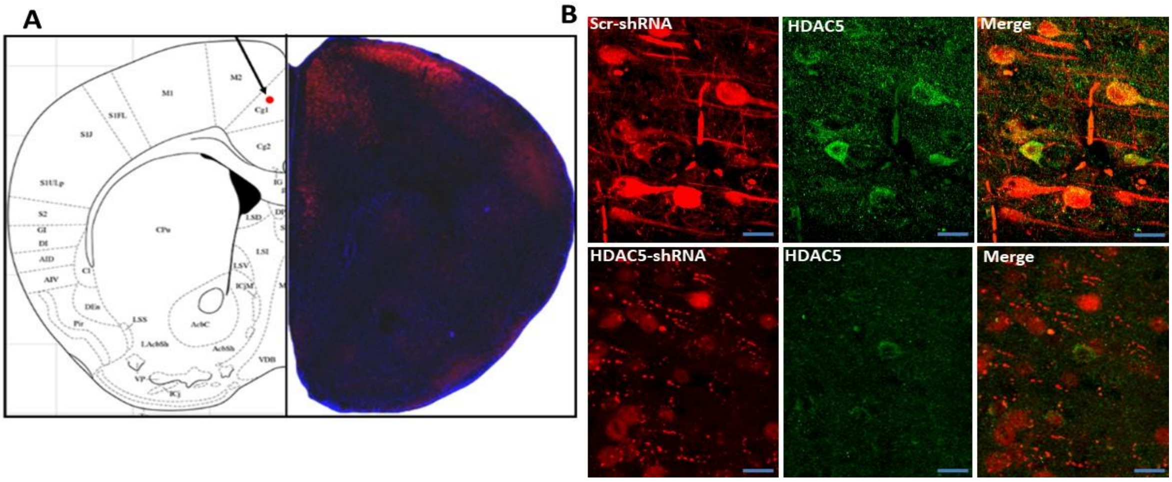


**Figure S9**. Stereotaxic injection of HDAC5-shRNA induced HDAC5 knockdown in the PFC. (**A**) Representative tile scan image showing the location of the AAV virus injection and expression of red fluorescence protein (RFP) in the PFC. (**B**) Representative immunostaining the PFC sections showing the colocalization of HDAC5 (green) staining with the RFP. A reduction in the HDAC5 expression was observed in the RFP-positive cells of the mice expressing HDAC5-shRNA compared to the mice expressing scrambled-shRNA (scr-shRNA). Scale bar= 20µm

### Vector map and sequence of HDAC-shRNA

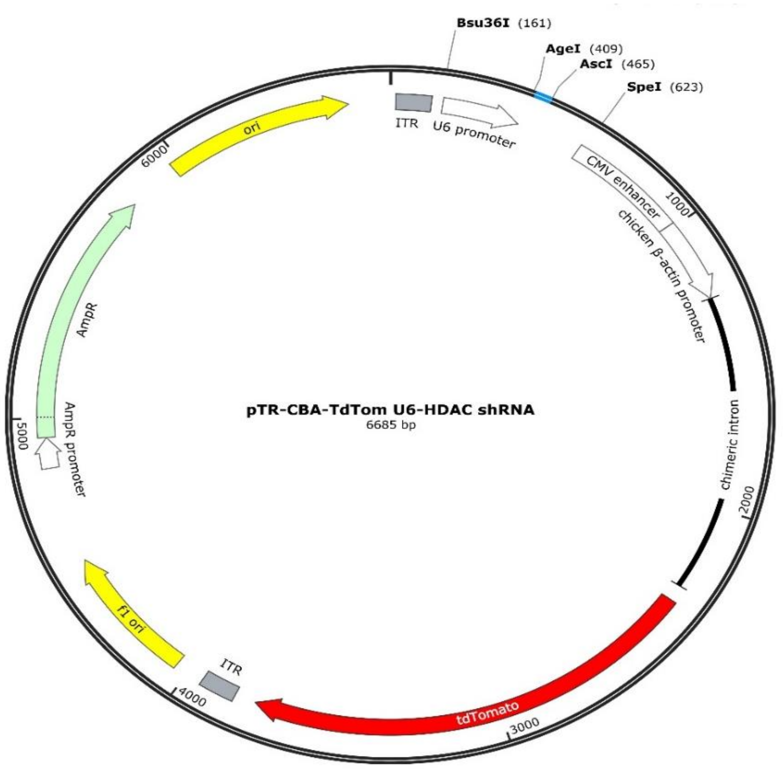


GGGGGGGGGGGGGGGGTTGGCCACTCCCTCTCTGCGCGCTCGCTCGCTCACTGAGGCCGGGCGACCA
AAGGTCGCCCGACGCCCGGGCTTTGCCCGGGCGGCCTCAGTGAGCGAGCGAGCGCGCAGAGAGGGAG
TGGCCAACTCCATCACTAGGGGTTCCTGAGGGCCTATTTCCCATGATTCCTTCATATTTGCATATACGATA
CAAGGCTGTTAGAGAGATAATTGGAATTAATTTGACTGTAAACACAAAGATATTAGTACAAAATACGTGA
CGTAGAAAGTAATAATTTCTTGGGTAGTTTGCAGTTTTAAAATTATGTTTTAAAATGGACTATCATATGCT
TACCGTAACTTGAAAGTATTTCGATTTCTTGGCTTTATATATCTTGTGGAAAGGACGAAACACCGGT**TGG**
**GCAAGATCCTTACCAACTCGAGTTGGTAAGGATCTTGCCCATTTTTG**GCGCGCCTCAATATTGGCCATTA
GCCATATTATTCATTGGTTATATAGCATAAATCAATATTGGATATTGGCCATTGCATACGTTGTATCTATAT
CATAATATGTACATTTATATTGGCTCATGTCCAATATGACCGCCATGTTGGCATTGATTATTGACTAGTTAT
TAATAGTAATCAATTACGGGGTCATTAGTTCATAGCCCATATATGGAGTTCCGCGTTACATAACTTACGGT
AAATGGCCCGCCTGGCTGACCGCCCAACGACCCCCGCCCATTGACGTCAATAATGACGTATGTTCCCATA
GTAACGCCAATAGGGACTTTCCATTGACGTCAATGGGTGGAGTATTTACGGTAAACTGCCCACTTGGCAG
TACATCAAGTGTATCATATGCCAAGTCCGCCCCCTATTGACGTCAATGACGGTAAATGGCCCGCCTGGCA
TTATGCCCAGTACATGACCTTACGGGACTTTCCTACTTGGCAGTACATCTACGTATTAGTCATCGCTATTA
CCATGGTCGAGGTGAGCCCCACGTTCTGCTTCACTCTCCCCATCTCCCCCCCCTCCCCACCCCCAATTTTGT
ATTTATTTATTTTTTAATTATTTTGTGCAGCGATGGGGGCGGGGGGGGGGGGGGGGCGCGCGCCAGGC
GGGGCGGGGCGGGGCGAGGGGCGGGGCGGGGCGAGGCGGAGAGGTGCGGCGGCAGCCAATCAGAG
CGGCGCGCTCCGAAAGTTTCCTTTTATGGCGAGGCGGCGGCGGCGGCGGCCCTATAAAAAGCGAAGCG
CGCGGCGGGCGGGAGTCGCTGCGACGCTGCCTTCGCCCCGTGCCCCGCTCCGCCGCCGCCTCGCGCCGC
CCGCCCCGGCTCTGACTGACCGCGTTACTCCCACAGGTGAGCGGGCGGGACGGCCCTTCTCCTCCGGGC
TGTAATTAGCGCTTGGTTTAATGACGGCTTGTTTCTTTTCTGTGGCTGCGTGAAAGCCTTGAGGGGCTCC
GGGAGGGCCCTTTGTGCGGGGGGGAGCGGCTCGGGGGGTGCGTGCGTGTGTGTGTGCGTGGGGAGCG
CCGCGTGCGGCCCGCGCTGCCCGGCGGCTGTGAGCGCTGCGGGCGCGGCGCGGGGCTTTGTGCGCTCC
GCAGTGTGCGCGAGGGGAGCGCGGCCGGGGGCGGTGCCCCGCGGTGCGGGGGGGGCTGCGAGGGGA
ACAAAGGCTGCGTGCGGGGTGTGTGCGTGGGGGGGTGAGCAGGGGGTATGGGCGCGGCGGTCGGGC
TGTAACCCCCCCCTGCACCCCCCTCCCCGAGTTGCTGAGCACGGCCCGGCTTCGGGTGCGGGGCTCCGTA
CGGGGCGTGGCGCGGGGCTCGCCGTGCCGGGCGGGGGGTGGCGGCAGGTGGGGGTGCCGGGCGGGG
CGGGGCCGCCTCGGGCCGGGGAGGGCTCGGGGGAGGGGCGCGGCGGCCCCCGGAGCGCCGGCGGCT

GTCGAGGCGCGGCGAGCCGCAGCCATTGCCTTTTATGGTAATCGTGCGAGAGGGCGCAGGGACTTACTT
TGTCCCAAATCTGTGCGGAGCCGAAATCTGGGAGGCGCCGCCGCACCCCCTCTAGCGGGCGCGGGGCG
AAGCGGTGCGGCGCCGGCAGGAAGGAAATGGGCGGGGAGGGCCTTCGTGCGTCGCCGCGCCGCCGTC
CCCTTCTCCCTCTCCAGCCTCGGGGCTGTCCGCGGGGGGACGGCTGCCTTCGGGGGGGACGGGGCAGG
GCGGGGTTCGGCTTCTGGCGTGTGACCGGCGGCTCTAGAGCCTCTGCTAACCATGTTCATGCCTTCTTCT
TTTTCCTACAGCTCCTGGGCAACGTGCTGGTTATTGTGCTGTCTCATCATTTTGGCAAAGAATTCATGGTG
AGCAAGGGCGAGGAGGTCATCAAAGAGTTCATGCGCTTCAAGGTGCGCATGGAGGGCTCCATGAACGG
CCACGAGTTCGAGATCGAGGGCGAGGGCGAGGGCCGCCCCTACGAGGGCACCCAGACCGCCAAGCTGA
AGGTGACCAAGGGCGGCCCCCTGCCCTTCGCCTGGGACATCCTGTCCCCCCAGTTCATGTACGGCTCCAA
GGCGTACGTGAAGCACCCCGCCGACATCCCCGATTACAAGAAGCTGTCCTTCCCCGAGGGCTTCAAGTG
GGAGCGCGTGATGAACTTCGAGGACGGCGGTCTGGTGACCGTGACCCAGGACTCCTCCCTGCAGGACG
GCACGCTGATCTACAAGGTGAAGATGCGCGGCACCAACTTCCCCCCCGACGGCCCCGTAATGCAGAAGA
AGACCATGGGCTGGGAGGCCTCCACCGAGCGCCTGTACCCCCGCGACGGCGTGCTGAAGGGCGAGATC
CACCAGGCCCTGAAGCTGAAGGACGGCGGCCACTACCTGGTGGAGTTCAAGACCATCTACATGGCCAAG
AAGCCCGTGCAACTGCCCGGCTACTACTACGTGGACACCAAGCTGGACATCACCTCCCACAACGAGGACT
ACACCATCGTGGAACAGTACGAGCGCTCCGAGGGCCGCCACCACCTGTTCCTGGGGCATGGCACCGGCA
GCACCGGCAGCGGCAGCTCCGGCACCGCCTCCTCCGAGGACAACAACATGGCCGTCATCAAAGAGTTCA
TGCGCTTCAAGGTGCGCATGGAGGGCTCCATGAACGGCCACGAGTTCGAGATCGAGGGCGAGGGCGAG
GGCCGCCCCTACGAGGGCACCCAGACCGCCAAGCTGAAGGTGACCAAGGGCGGCCCCCTGCCCTTCGCC
TGGGACATCCTGTCCCCCCAGTTCATGTACGGCTCCAAGGCGTACGTGAAGCACCCCGCCGACATCCCCG
ATTACAAGAAGCTGTCCTTCCCCGAGGGCTTCAAGTGGGAGCGCGTGATGAACTTCGAGGACGGCGGTC
TGGTGACCGTGACCCAGGACTCCTCCCTGCAGGACGGCACGCTGATCTACAAGGTGAAGATGCGCGGCA
CCAACTTCCCCCCCGACGGCCCCGTAATGCAGAAGAAGACCATGGGCTGGGAGGCCTCCACCGAGCGCC
TGTACCCCCGCGACGGCGTGCTGAAGGGCGAGATCCACCAGGCCCTGAAGCTGAAGGACGGCGGCCAC
TACCTGGTGGAGTTCAAGACCATCTACATGGCCAAGAAGCCCGTGCAACTGCCCGGCTACTACTACGTG
GACACCAAGCTGGACATCACCTCCCACAACGAGGACTACACCATCGTGGAACAGTACGAGCGCTCCGAG
GGCCGCCACCACCTGTTCCTGTACGGCATGGACGAGCTGTACAAGTAGGCGGCCGCGACTCTAAATCGA
TAAGGATCTAGGAACCCCTAGTGATGGAGTTGGCCACTCCCTCTCTGCGCGCTCGCTCGCTCACTGAGGC
CGCCCGGGCAAAGCCCGGGCGTCGGGCGACCTTTGGTCGCCCGGCCTCAGTGAGCGAGCGAGCGCGCA
GAGAGGGAGTGGCCAACCCCCCCCCCCCCCCCCTGCAGCCTGGCGTAATAGCGAAGAGGCCCGCACCG
ATCGCCCTTCCCAACAGTTGCGTAGCCTGAATGGCGAATGGCGCGACGCGCCCTGTAGCGGCGCATTAA
GCGCGGCGGGTGTGGTGGTTACGCGCAGCGTGACCGCTACACTTGCCAGCGCCCTAGCGCCCGCTCCTT
TCGCTTTCTTCCCTTCCTTTCTCGCCACGTTCGCCGGCTTTCCCCGTCAAGCTCTAAATCGGGGGCTCCCTT
TAGGGTTCCGATTTAGTGCTTTACGGCACCTCGACCCCAAAAAACTTGATTAGGGTGATGGTTCACGTAG
TGGGCCATCGCCCTGATAGACGGTTTTTCGCCCTTTGACGTTGGAGTCCACGTTCTTTAATAGTGGACTCT
TGTTCCAAACTGGAACAACACTCAACCCTATCTCGGTCTATTCTTTTGATTTATAAGGGATTTTGCCGATTT
CGGCCTATTGGTTAAAAAATGAGCTGATTTAACAAAAATTTAACGCGAATTTTAACAAAATATTAACGTTT
ACAATTTCCTGATGCGCTATTTTCTCCTTACGCATCTGTGCGGTATTTCACACCGCATATGGTGCACTCTCA
GTACAATCTGCTCTGATGCCGCATAGTTAAGCCAGCCCCGACACCCGCCAACACCCGCTGACGCGCCCTG
ACGGGCTTGTCTGCTCCCGGCATCCGCTTACAGACAAGCTGTGACCGTCTCCGGGAGCTGCATGTGTCAG
AGGTTTTCACCGTCATCACCGAAACGCGCGAGACGAAAGGGCCTCGTGATACGCCTATTTTTATAGGTTA
ATGTCATGATAATAATGGTTTCTTAGACGTCAGGTGGCACTTTTCGGGGAAATGTGCGCGGAACCCCTAT
TTGTTTATTTTTCTAAATACTTTCAAATATGTATCCGCTCATGAGACAATAACCCTGATAAATGCTTCAATA
ATATTGAAAAAGGAAGAGTATGAGTATTCAACATTTCCGTGTCGCCCTTATTCCCTTTTTTGCGGCATTTT
GCCTTCCTGTTTTTGCTCACCCAGAAACGCTGGTGAAAGTAAAAGATGCTGAAGATCAGTTGGGTGCACG
AGTGGGTTACATCGAACTGGATCTCAACAGCGGTAAGATCCTTGAGAGTTTTCGCCCCGAAGAACGTTTT
CCAATGATGAGCACTTTTAAAGTTCTGCTATGTGGCGCGGTATTATCCCGTATTGACGCCGGGCAAGAGC
AACTCGGTCGCCGCATACACTATTCTCAGAATGACTTGGTTGAGTACTCACCAGTCACAGAAAAGCATCT
TACGGATGGCATGACAGTAAGAGAATTATGCAGTGCTGCCATAACCATGAGTGATAACACTGCGGCCAA

CTTACTTCTGACAACGATCGGAGGACCGAAGGAGCTAACCGCTTTTTTGCACAACATGGGGGATCATGTA
ACTCGCCTTGATCGTTGGGAACCGGAGCTGAATGAAGCCATACCAAACGACGAGCGTGACACCACGATG
CCTGTAGCAATGGCAACAACGTTGCGCAAACTATTAACTGGCGAACTACTTACTCTAGCTTCCCGGCAAC
AATTAATAGACTGGATGGAGGCGGATAAAGTTGCAGGACCACTTCTGCGCTCGGCCCTTCCGGCTGGCT
GGTTTATTGCGGATAAATCTGGAGCCGGTGAGCGTGGGTCTCGCGGTATCATTGCAGCACTGGGGCCAG
ATGGTAAGCCCTCCCGTATCGTAGTTATCTACACGACGGGGAGTCAGGCAACTATGGATGAACGAAATA
GACAGATCGCTGAGATAGGTGCCTCACTGATTAAGCATTGGTAACTGTCAGACCAAGTTTACTCATATAT
ACTTTAGATTGATTTAAAACTTCATTTTTAATTTAAAAGGATCTAGGTGAAGATCCTTTTTGATAATCTCAT
GACCAAAATCCCTTAACGTGAGTTTTCGTTCCACTGAGCGTCAGACCCCGTAGAAAAGATCAAAGGATCT
TCTTGAGATCCTTTTTTTCTGCGCGTAATCTGCTGCTTGCAAACAAAAAAACCACCGCTACCAGCGGTGGT
TTGTTTGCCGGATCAAGAGCTACCAACTCTTTTTCCGAAGGTAACTGGCTTCAGCAGAGCGCAGATACCA
AATACTGTCCTTCTAGTGTAGCCGTAGTTAGGCCACCACTTCAAGAACTCTGTAGCACCGCCTACATACCT
CGCTCTGCTAATCCTGTTACCAGTGGCTGCTGCCAGTGGCGATAAGTCGTGTCTTACCGGGTTGGACTCA
AGACGATAGTTACCGGATAAGGCGCAGCGGTCGGGCTGAACGGGGGGTTCGTGCACACAGCCCAGCTT
GGAGCGAACGACCTACACCGAACTGAGATACCTACAGCGTGAGCATTGAGAAAGCGCCACGCTTCCCGA
AGGGAGAAAGGCGGACAGGTATCCGGTAAGCGGCAGGGTCGGAACAGGAGAGCGCACGAGGGAGCT
TCCAGGGGGAAACGCCTGGTATCTTTATAGTCCTGTCGGGTTTCGCCACCTCTGACTTGAGCGTCGATTTT
TGTGATGCTCGTCAGGGGGGCGGAGCCTATGGAAAAACGCCAGCAACGCGGCCTTTTTACGGTTCCTGG
CCTTTTGCTGGCCTTTTGCTCACATGTTCTTTCCTGCGTTATCCCCTGATTCTGTGGATAACCGTATTACCG
CCTTTGAGTGAGCTGATACCGCT
